# Holosteans contextualize the role of the teleost genome duplication in promoting the rise of evolutionary novelties in the ray-finned fish innate immune system

**DOI:** 10.1101/2021.06.11.448072

**Authors:** Alex Dornburg, Dustin J. Wcisel, Katerina Zapfe, Emma Ferraro, Lindsay Roupe-Abrams, Andrew W. Thompson, Ingo Braasch, Tatsuya Ota, Jeffrey A. Yoder

## Abstract

Over 99% of ray-finned fishes (Actinopterygii) are teleosts, a clade that comprises half of all living vertebrates that have diversified across virtually all fresh and saltwater ecosystems. This ecological diversity raises the question of how the immunogenetic diversity required to persist under heterogeneous pathogen pressures evolved. The teleost genome duplication (TGD) has been hypothesized as the evolutionary event that provided the genomic substrate for rapid genomic evolution and innovation. However, studies of putative teleost-specific innate immune receptors have been largely limited to comparisons either among teleosts or between teleosts and distantly related vertebrate clades such as tetrapods. Here we describe and characterize the receptor diversity of two clustered innate immune gene families in the teleost sister lineage: Holostei (bowfin and gars). Using genomic and transcriptomic data, we provide a detailed investigation of the phylogenetic history and conserved synteny of gene clusters encoding diverse immunoglobulin domain-containing proteins (DICPs) and novel immune-type receptors (NITRs). These data demonstrate an ancient linkage of DICPs to the major histocompatibility complex (MHC) and reveal an evolutionary origin of NITR variable-joining (VJ) exons that predate the TGD by at least 50 million years. Further characterizing the receptor diversity of Holostean DICPs and NITRs illuminates a sequence diversity that rivals the diversity of these innate immune receptor families in many teleosts. Taken together, our findings provide important historical context for the evolution of these gene families that challenge prevailing expectations concerning the consequences of the TGD during actinopterygiian evolution.

## Introduction

With over 33,000 species, teleost fishes are the most diverse clade of living ray-finned fishes (Actinopterygii). A hallmark of their diversification history is the ability to repeatedly colonize and radiate across virtually all of the planet’s fresh and saltwater ecosystems (Seehausen 2006; Friedman et al. 2013; Davis et al. 2014; Price et al. 2014; Brawand et al. 2014; Dornburg et al. 2017; Salzburger 2018; Daane et al. 2019). While numerous studies have identified mechanisms that underlie teleost morphological and species diversification dynamics (Price et al. 2014; Sibert et al. 2018; Iglesias et al. 2018; Gajdzik et al. 2019; Near and Kim 2021), the role of the teleost genome duplication (TGD) (Braasch and Postlethwait 2012; Glasauer and Neuhauss 2014) event as a substrate that promoted the evolution of immunogenetic novelty in the early evolutionary history of teleosts remains unclear. On the one hand, investigations of teleost genomes have resulted in descriptions of several families of innate immune receptors putatively unique to teleosts (Yoder and Litman 2011; Montgomery et al. 2011; Rodríguez-Nunez et al. 2014; Wcisel and Yoder 2016; Traver and Yoder 2020). On the other, with very few exceptions (Boudinot et al. 2014; Conant 2020) most comparative immunogenetic studies have been limited to investigations among teleosts (Ferraresso et al. 2009; Rebl et al. 2010; Montgomery et al. 2011; Aoki et al. 2013; Pietretti and Wiegertjes 2014; Rodríguez-Nunez et al. 2014; Wcisel and Yoder 2016) or between teleosts and distantly related vertebrate clades such as tetrapods (Yoder and Litman 2011; Langevin et al. 2013; Kasahara and Flajnik 2019). These studies are of incredible value, but do not consider the potential sequence diversity within the few living non-teleost actinopterygians. Such a comparison is critical if we are to disentangle teleost-specific patterns of evolution from the overall sequence diversity and genomic architecture of the general ray-finned fish immune system.

The closest relatives to living teleosts are holosteans (Grande 2010; Near et al. 2012b; Braasch et al. 2016; Betancur-R et al. 2017; Wcisel et al. 2020; Hughes et al. 2018), vestiges of two clades of ancient fishes that flourished during the Mesozoic (Smithwick and Stubbs 2018). Recent sequencing of both the spotted gar (*Lepisosteus oculatus*) (Braasch et al. 2016) and the bowfin (*Amia calva*) genomes (Thompson et al. 2021) found that portions of the genomes of these fishes demonstrate surprising conservation with elements from tetrapods, thereby revealing likely characteristics of early bony vertebrate genomes. For example, the bowfin genome revealed an extensive linkage of MHC class I, class II and class III loci (Thompson et al. 2021), a condition found in humans but not observed in any teleost fish and unresolved in gar (Braasch et al. 2016; Yamaguchi and Dijkstra 2019). Moreover, clusters of diverse immunoglobulin-domain containing proteins (DICPs) are encoded within the MHC locus of bowfin. DICPs have been identified in teleosts, holosteans, and coelacanth and possess one or more extracellular Ig domains that fall into two types termed D1 and D2 (Haire et al. 2012; Boudinot et al. 2014; Rodriguez-Nunez et al. 2016; Wcisel and Yoder 2016; Gao et al. 2018). In zebrafish, DICPs are expressed by lymphocytes and myeloid cells, are predicted to play important roles in pathogen recognition, and may regulate interferon signaling (Haire et al. 2012; Carmona et al. 2017; Gao et al. 2018). In bowfin, these multigene clusters are surrounded by well-conserved genes whose synteny can be traced to both teleost (zebrafish) and tetrapod (human) genomes (Thompson et al. 2021). As such, investigating the evolutionary history of these gene clusters may help illuminate core motifs of the ancestral, (‘proto’ or ‘Ur’), vertebrate MHC (Abi Rached et al. 1999; Kasahara 1999).

In addition to loci associated with the MHC locus, the spotted gar genome revealed the first evidence for the existence of novel immune-type receptors (NITRs) outside of teleosts (Braasch et al. 2016; Wcisel et al. 2017). NITR clusters have been identified in all examined teleost lineages (Strong et al. 1999; Yoder et al. 2004; Desai et al. 2008; Yoder 2009; Ferraresso et al. 2009; Wcisel and Yoder 2016) and are predicted to function as natural killer cell receptors (Litman et al. 2001; Yoder et al. 2010). Typically, NITRs possess two extracellular Ig domains, one membrane-distal variable (V)-like domain and one membrane-proximal intermediate (I)-like domain. The majority of NITR V and I domains include a C-terminal joining (J)-like sequence. Given that the V-J and I-J sequences are encoded in single exons with no evidence for recombination, this raises the possibility that NITRs are derived from the progenitor receptor sequences that gave rise to VJ and subsequently VDJ recombination in Immunoglobulin (Ig) and T-Cell Receptor (TCR) genes of the adaptive immune system of jawed vertebrates (Yoder 2009). In the absence of a detailed analysis of holostean NITRs, it remains unclear whether the origin of their diversity aligns with or predates the origin of teleosts and the TGD.

NITRs and DICPs also share similarities beyond possessing extracellular Ig domains. An additional unifying feature of the DICP and NITR families is the inclusion of inhibitory, activating, secreted and functionally ambiguous forms (Wcisel and Yoder 2016). Inhibitory DICPs and NITRs encode one or more cytoplasmic immunoreceptor tyrosine-based inhibition motif (ITIM; S/I/V/LxYxxI/V/L) or ITIM-like sequences (itim), whereas activating receptors possess an intramembranous charged residue permitting association with activating adaptor proteins (e.g. Dap12) (Wei et al. 2007; Wcisel and Yoder 2016). Both receptor families also include secreted forms that lack a transmembrane domain and functionally ambiguous forms that possess a transmembrane domain, but lack any identifiable signaling motif. However, whether all of these different functional forms are represented in holosteans or if some are restricted to teleosts remains to be determined.

Here we integrate phylogenetic, genomic, and transcriptomic analyses of the spotted gar and bowfin DICPs and NITRs. We begin by providing the first description of the diversity of NITRs and DICPs across holosteans, illuminating evidence for functionally analogous sequences between the deeply divergent lineages of gar and bowfin. We then use phylogenetic approaches to estimate the evolutionary history of each receptor family across the common neopterygian fish ancestor of holosteans and teleosts. This allowed us to test whether the TGD is correlated with higher receptor diversity in teleosts versus equivalent signatures of diversification within holosteans. We then assess patterns of conserved synteny between holostean receptor families, allowing us to assess the degree to which gene regions surrounding holosteans are syntenic to those of teleosts. Collectively, our findings provide the historical framework necessary to contextualize changes of these receptor families in teleosts and neopteryigans. More broadly, these findings provide a new perspective for future work concerning the consequences of the TGD on teleost fish immunogenetic receptor diversity.

## Methods

### Overview of the bowfin genome

Our analysis is based on the recently published bowfin genome that contains 1,958 scaffolds and 23 pseudochromosomes (named Aca_scaf_1 through Aca_scaf_23) that contain 99% of the assembly and match bowfin’s chromosome number (Thompson et al. 2021). These pseudochromosomes are consistent with a chromosome-level genome assembly, but have not yet been definitively assigned to individual chromosomes. As such, we retain the Aca_scaf nomenclature in this manuscript. Genes in the bowfin genome were annotated using the MAKER genome annotation software (Holt and Yandell 2011), reporting a total of approximately 22,000 protein-coding genes (Thompson et al. 2021). This gene annotation applied the prefixes AMCG (AMia Calva Gene) and AMCP (AMia Calva Protein). We refined these predictions in our delimitation of bowfin DICP and NITR sequences.

### Bowfin DICP and NITR nomenclature

Bowfin DICP and NITR sequences were named consecutively: *dicp1, dicp2, dicp3* etc., and *nitr1, nitr2, nitr3* etc. Putative pseudogenes were assigned the next number in the symbol series and suffixed by a “p”. Sequences were considered pseudogenes if a predicted exon possessed an internal stop codon or if a solitary exon was identified. It is emphasized that due to recent lineage-specific diversification of the DICP and NITR families, one-to-one genetic orthologs are not identifiable between species, and gene names reflect only the order in which they were identified. For example medaka *NITR1a* is not a “true” ortholog of either zebrafish *nitr1a* or pufferfish *NITR1* (Desai et al. 2008).

### Identification of bowfin DICP genes and transcripts

The identification and organization of bowfin genes *dicp1* through *dicp20* on Aca_scaf_14 was reported previously (Thompson et al. 2021). Note that the genome sequence of the *dicp18p* pseudogene can only encode a partial Ig domain (32 residues) (Thompson et al. 2021) for which additional analyses are not possible. As such, we excluded this gene from all analyses below. tBLASTn searches of the bowfin reference genome were conducted using spotted gar DICP Ig domain sequences as queries to search for additional DICPs in other regions of the bowfin genome. BLAST searches of publicly available bowfin RNA-seq (**Supplementary Note 1**) were used to identify DICP transcripts. Specifically, tBLASTn searches (E value cutoff E-6) of bowfin transcript sequences available through the PhyloFish database (http://phylofish.sigenae.org/) (Pasquier et al. 2016) and bowfin transcriptome data from the immune tissues of a single adult fish (bowfin 0039) (Thompson et al. 2021) were conducted using spotted gar DICP Ig domain sequences as queries. Syntenic relationships between bowfin and gar were determined using reciprocal BLASTp searches against the spotted gar genome. To ensure genes with multiple isoforms were only represented by one protein sequence, only the first protein sequence encountered in the annotation file for each genome was used in the BLAST analysis. The results of the BLAST searches were subjected to collinear analyses using MCScanX (Wang et al. 2012).

### Identification of bowfin NITR genes and transcripts

In our initial search of the bowfin genome-predicted proteins (e.g. AMCP sequences) for NITR sequences, we employed spotted gar NITR I domains (Wcisel et al. 2017) as queries for BLASTP searches (e value cutoff < 1e-15) as the high rate of sequence evolution in NITR V domains limits their utility for effective homology searches over deep evolutionary time scales. In order to identify bowfin NITR transcripts, we used the candidate NITR sequences encoded by the genes described above and all reported spotted gar NITR I domains (Wcisel et al. 2017) as queries for tBLASTn searches (e values <1e-10) against immune tissue RNA-seq from bowfin 0039 (Thompson et al. 2021) and from bowfin available from PhyloFish (Pasquier et al. 2016) (**Supplementary Note 1**). Resultant protein sequences were manually inspected and those lacking an Ig domain or found to encode Ig light chain proteins were excluded. Remaining sequences were trimmed to remove untranslated sequences.

### Reanalyses of the bowfin NITR cluster

As our pilot analysis indicated the presence of multiple I domains in individual predicted bowfin AMCP proteins (which would be an unprecedented NITR protein architecture), we reanalyzed the delimitation of bowfin NITRs using all identified bowfin I domain peptide sequences as queries for tBLASTn searches (e values <1e-10) of the reference genome. Sequences that did not possess at least four of the six conserved cysteines within the I domain were removed. We replicated these tBLASTn searches to identify additional NITR V domains, but restricted sequences to those with a corresponding I domain in close genomic proximity. We combined all candidate NITR nucleotide sequences from the genomic and transcriptomic searches into Splign (Kapustin et al. 2008) to map all possible exons from genes and transcripts. This allowed us to manually link adjacent NITR exons into individual NITR genes and create a genomic map of the NITR gene cluster. Exonic maps were generated using the ggplot 2 package (Wickham 2011) in R 4.0.2. Syntenic relationships were determined as for DICPs.

### Protein sequence analyses

Protein domains were identified using SMART (Letunic and Bork 2018) and SignalP (Almagro Armenteros et al. 2019). Protein sequences from bowfin were compared to DICP or NITR sequences from spotted gar (Wcisel et al. 2017) and representative teleosts in which these receptors had previously been described [zebrafish (*Danio rerio*), carp (*Cyprinus carpio* and *Ctenopharyngodon idella*), pufferfish (*Takifugu rubripes* and *Sphoeroides nephelus*), salmon (*Salmo salar*), medaka (*Oryzias latipes)*, and tilapia (*Oreochromis niloticus*) (Strong et al. 1999; Yoder et al. 2004, 2008; Desai et al. 2008; Haire et al. 2012; Rodriguez-Nunez et al. 2016)]. In addition, a candidate DICP from coelacanth (*Latimeria chalumnae*) was included (Boudinot et al. 2014). Sequences were aligned using Clustal Omega (Sievers and Higgins 2018) or MAFFT (Katoh and Standley 2014; Nakamura et al. 2018). Boxshade (version 3.21) alignment plots were made using the MAFFT-based alignment and manually annotated (https://embnet.vital-it.ch/software/BOX_form.html).

To determine the best-fit model of amino-acid substitution and infer a maximum likelihood phylogeny of DICP and NITR Ig domains, the alignment of each sequence was analyzed using IQ-TREE (Nguyen et al. 2015; Kalyaanamoorthy et al. 2017). The candidate pool of substitution models spanned all common amino acid exchange rate matrices (e.g., JTT (Jones et al. 1992), WAG (Whelan and Goldman 2001) to protein mixture models such as empirical profile mixture models (Quang et al. 2008), and also included parameters to accommodate among site rate variation (e.g., discrete gamma (Yang 1994) or a free rate model (Soubrier et al. 2012). The best-fit substitution model for each alignment was selected using Akaike information criterion in IQ-TREE (Nguyen et al. 2015; Kalyaanamoorthy et al. 2017). Node support was assessed via 1,000 ultrafast bootstrap replicates (Minh et al. 2013; Hoang et al. 2018).

## Results and Discussion

### Bowfin DICPs are embedded within the MHC

We recently reported two DICP gene ‘clusters’ on bowfin pseudochromosome Aca_scaf_14 (**Fig. 1a**) (Thompson et al. 2021) and our blast searches of the bowfin genome identified an additional pseudogene *dicp21p* on pseudochromosome Aca_scaf_11 with no support for synteny with known DICPs in other species. We previously reported multiple DICP sequences from the spotted gar genome that are encoded on multiple genomic scaffolds (Wcisel et al. 2017) that our results demonstrate to share synteny with the Aca_scaf_14 DICP cluster (**Fig. 1b**) suggesting they might be linked in gar as well. We report twenty different bowfin DICP genes and pseudogenes (*dicp1* - *dicp20*) encoded in a cluster within the extended MHC region (that includes MHC class I, class II and class III genes) on Aca_scaf_14 (**Fig. 1a-b** and **Supplementary Table S1**). This linkage of DICPs to the MHC in bowfin sheds light on our previous finding that zebrafish DICPs are tightly linked to MHC class I genes (Rodriguez-Nunez et al. 2016). These results in bowfin reveal that it is likely that in early diverging ray-finned fishes, DICPs were encoded within an extended MHC region (including class I, II, and III genes). Following the TGD, this linkage was likely fragmented through the heterogeneous maintenance of duplicated MHC gene clusters on different chromosomes.

**Fig. 1.**
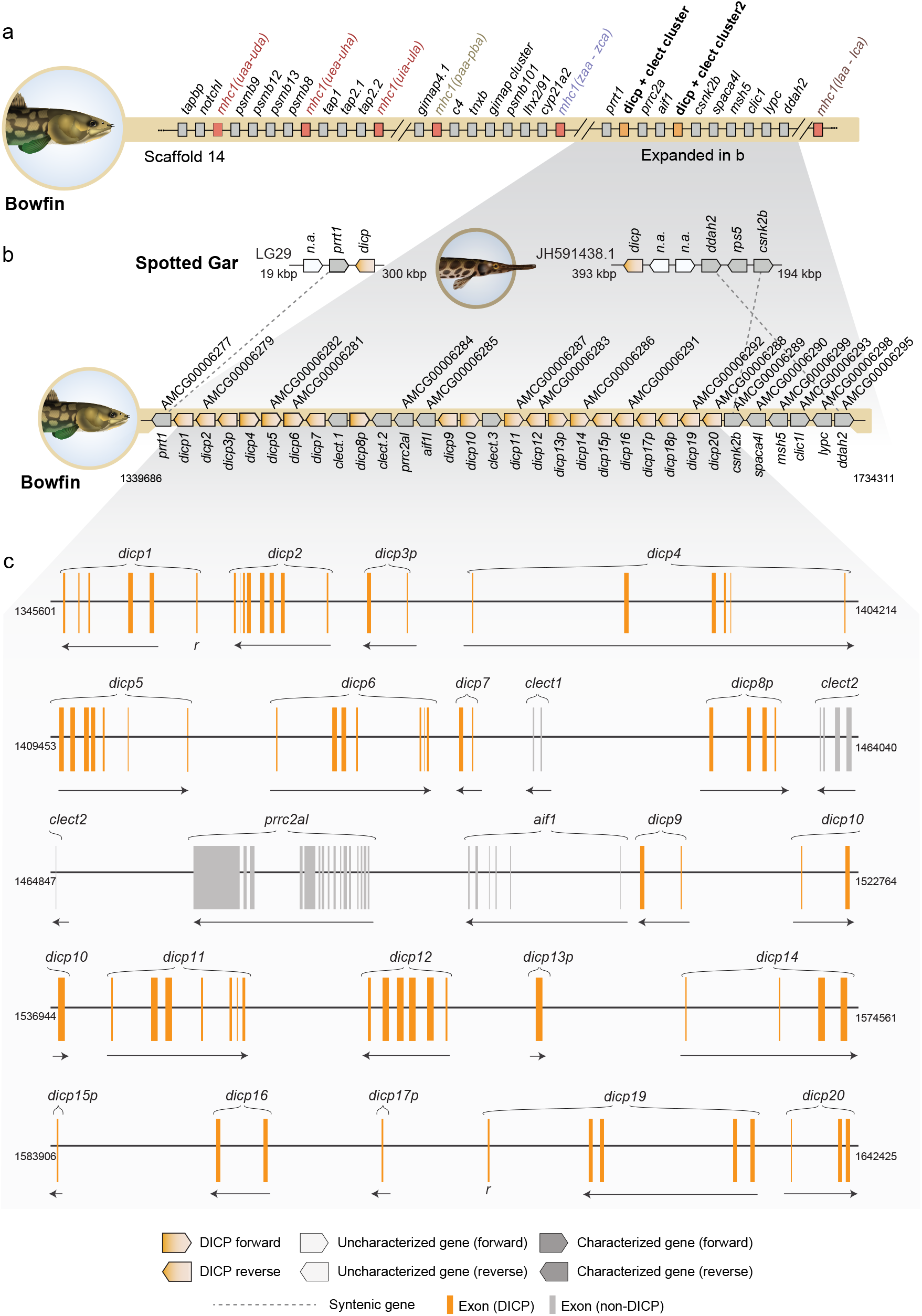
Bowfin DICPs are encoded within the MHC. **a** The main bowfin DICP gene cluster is encoded within the MHC on pseudochromosome Aca_scaf_14 (Thompson et al. 2021). The DICP gene cluster is flanked by clusters of MHC Class I Z lineage genes (*mhc1zaa, mhc1zba* and *mhc1zca*) and MHC Class I L lineages genes (*mhc1laa, mhc1lba* and *mhc1lca*). A single DICP pseudogene (*dicp21p*) is encoded on scaffold Aca_scaf_11 (not shown, see **Supplementary Table S1**). **b** The spotted gar MHC and DICP cluster are currently fragmented in the reference genome leading to limited, but compelling, evidence for conserved synteny between bowfin and gar (e.g. *prrt1, ddah2* and *csnk2b*). Each pentagon reflects a single gene with transcriptional orientation indicated. Gene sequence identifiers are shown above the genes (**a-b**) and DICP gene names are shown below the gene (**b**). **c** A detailed exon map represents the current knowledge of DICP genomic organization within the reference genome. Nucleotide position within Aca_scaf_14 is indicated on the left and right of each genomic region. T=predicted exons with reversed transcriptional orientation.

Teleost DICP genes typically encode at least five exons with the first four encoding a signal peptide sequence, a D1-type Ig domain, a D2-type Ig domain, and a transmembrane domain with subsequent exons encoding a cytoplasmic tail, although a few DICP genes encode four extracellular domains with a D1a-D2a-D1b-D2b organization (Haire et al. 2012). Our results demonstrate that many bowfin DICPs encode a similar number of exons as teleosts (**Fig. 1c** and **Supplementary Table S1)**. Although not all bowfin DICP genes can be confirmed to encode functional proteins (e.g. *dicp15p* and *dicp17p* were each identified as encoding a single transmembrane exon), at least eleven bowfin genes (*dicp2, dicp4, dicp5, dicp6, dicp10, dicp11, dicp12, dicp14, dicp16, dicp19* and *dicp20*) are predicted to encode functional DICPs (see below).

### Bowfin DICP Ig domain architectures mirror those of teleosts

The majority of DICPs in teleosts and spotted gar have been characterized by the presence of four conserved cysteines in their extracellular Ig domains, D1 and D2 (Haire et al. 2012; Rodriguez-Nunez et al. 2016; Wcisel et al. 2017; Gao et al. 2018). Two of these cysteines [C^23^ and C^104^, numbering based on the IMGT system (Lefranc et al. 2015)] likely play roles in stabilizing each Ig-fold (Williams and Barclay 1988). It has been suggested that the other, DICP-specific cysteines might promote DICP dimerization (Rodríguez-Nunez et al. 2014; Wcisel et al. 2017). Consistent with this expectation, we find that *dicp6, dicp10, dicp11, dicp14, dicp16* and *dicp20*, which likely encode proteins with a D1-D2 structure, possess all four conserved cysteines (**Figs. 2** and **3**). In contrast, *dicp2, dicp5, dicp12* and *dicp19*, which are predicted to encode proteins with a D1a-D2a-D1b-D2b structure, possess C^23^ and C^104^, but lack the majority of the DICP-specific cysteines. This subtle difference may indicate D1-D2 DICPs may be more likely to dimerize than D1a-D2a-D1b-D2b DICPs. Phylogenetic analyses of the bowfin DICP D1 and D2 domains support monophyly for the majority of the D1b and D2b domains (**Fig. 4**). This suggests a single early origin of the D1a-D2a-D1b-D2b motif with a subsequent loss of the D1b-D2b architecture in DICPs such as *dicp6, dicp9*, and *dicp11*. A possible mechanism for such an origin could be the loss of a transcription terminator signal from the upstream gene and the loss of a signal peptide from an adjacent downstream gene. Such a loss, along with alternative mRNA splicing, could facilitate the transcription of additional extracellular Ig domains in a single gene. Although speculative, such an event could also explain the finding of a D1a-D2a-D1b-D2b structure in zebrafish. The presence of this duplicated domain structure in bowfin suggests that repeating domain organization such as the D1a-D2a-D1b-D2b structure can arise from the clustered nature of gene families, rather than relying on major evolutionary events such as the TGD to provide a substrate for innovation.

**Fig. 2.**
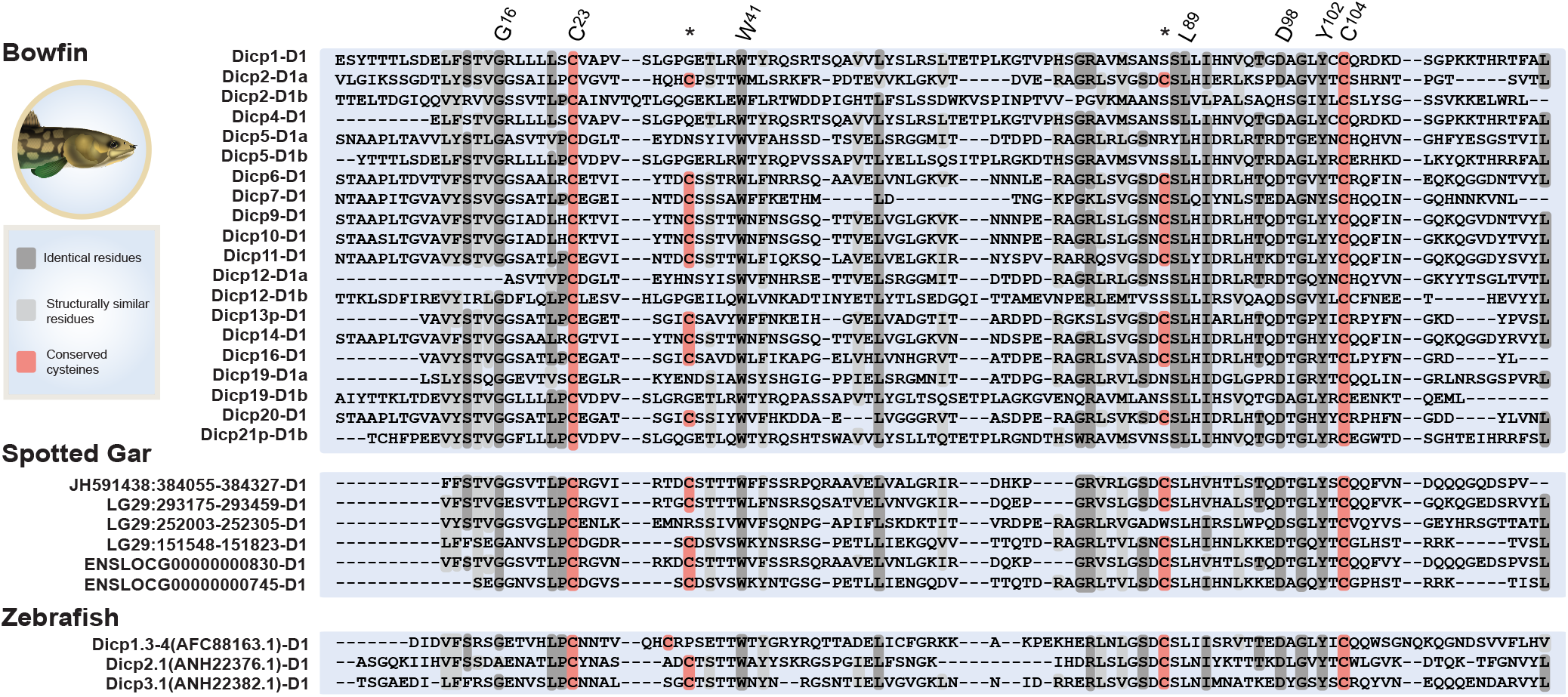
Conservation of the DICP D1 domain in bowfin. Protein sequence alignment of the D1 region of Bowfin DICPs compared with representative spotted gar and zebrafish sequences. Sequences are shaded by similarity with dashes indicating gaps in the alignment. Identical residues are shaded dark gray; structurally similar residues shaded light gray; and conserved cysteines shaded red. Key residues are numbered using the IMGT numbering system (Lefranc et al. 2015). Asterisks (*) indicate cysteines indicative of a DICP D1.

**Fig. 3.**
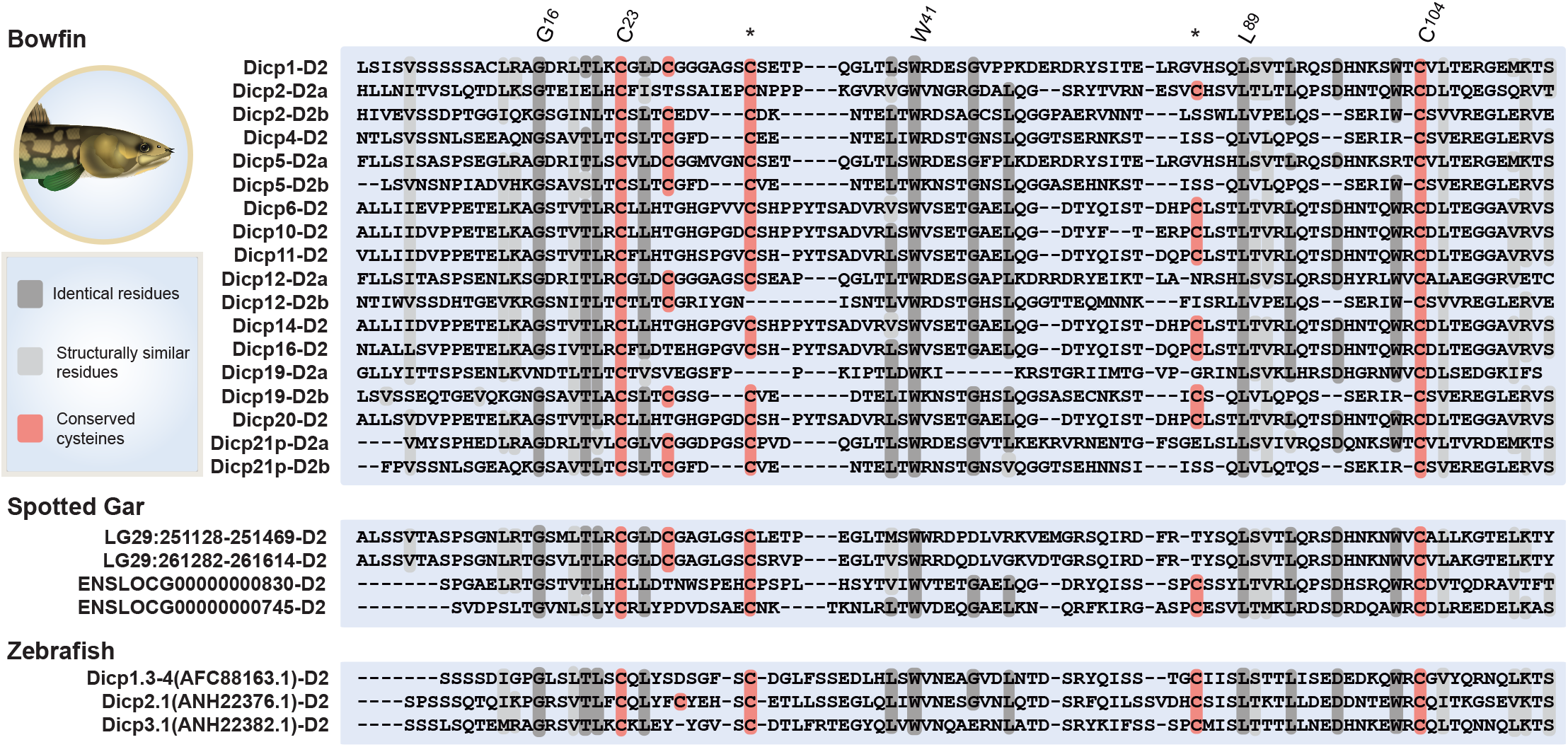
Conservation of the DICP D2 domain in bowfin. Protein sequence alignment of the D2 region of Bowfin DICPs compared with representative spotted gar and zebrafish sequences. Sequences are shaded by similarity with dashes indicating gaps in the alignment. Identical residues are shaded dark gray; structurally similar residues shaded light gray; and conserved cysteines shaded red. Key residues are numbered using the IMGT numbering system (Lefranc et al. 2015). Asterisks (*) indicate cysteines indicative of a DICP D2.

**Fig. 4.**
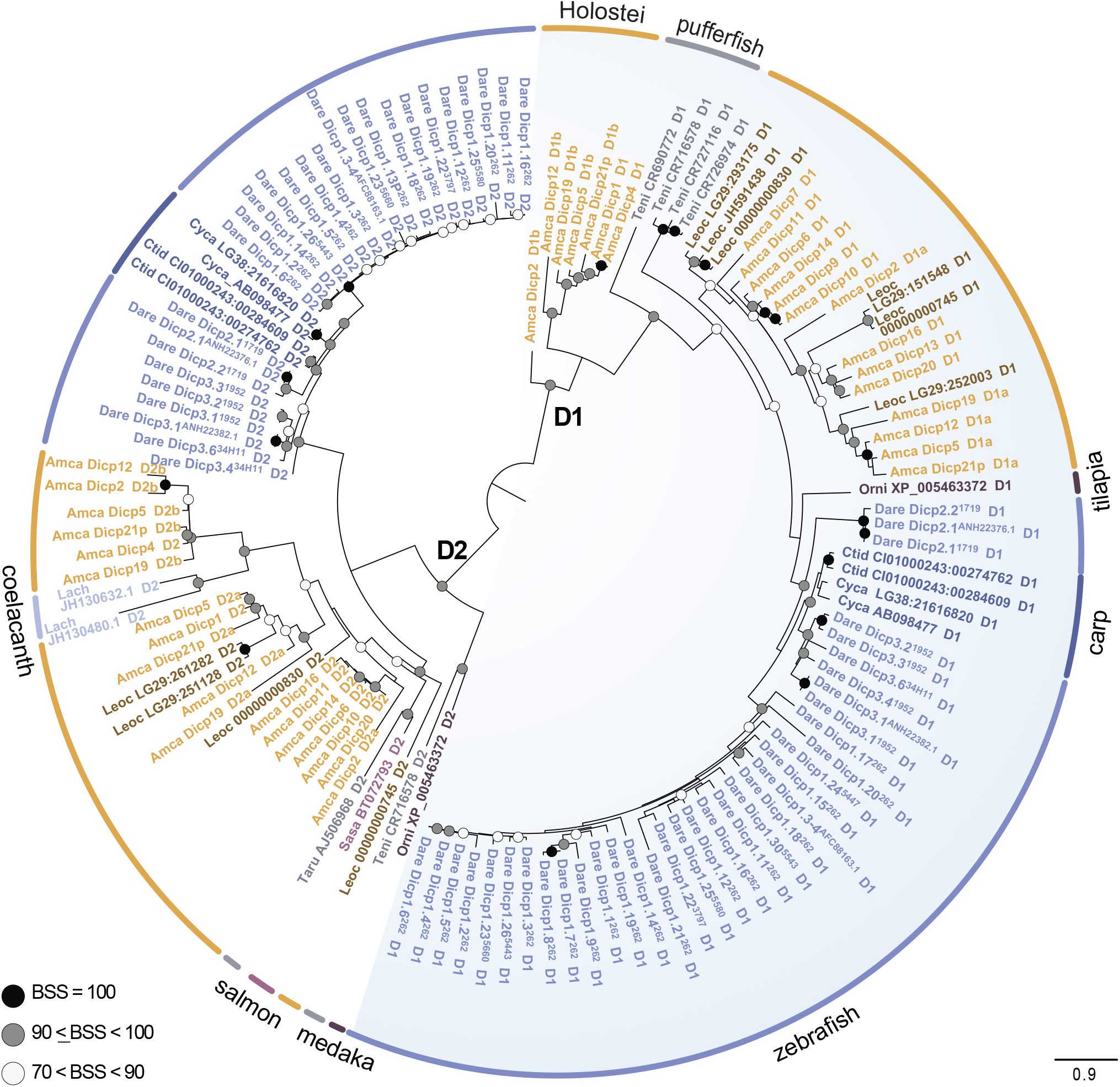
DICP diversity does not reflect holostean monophyly. Maximum likelihood phylogeny of DICP D1 and D2 domains inferred using IQ-TREE. Circles at nodes indicate bootstrap support values (BSS) with filled black circles black indicating BSS=100, gray circles indicating BSS values equal to or greater than 90 but less than 100, and white circles indicating BSS values greater than 70 but less than 90. Lineages are indicated by the color coded arcs surrounding the tree and colored taxon labels (light blue = zebrafish [*Danio rerio*, Dare]; dark blue = carp [*Cyprinus carpio*, Cyca and *Ctenopharyngodon idella*, Ctid]; turquoise = coelacanth [*Latimeria chalumnae*, Lach]; gray = pufferfish [*Tetraodon nigroviridis*, Teni]; dark purple = tilapia [*Oreochromis niloticus*, Orni]; light purple = salmon [*Salmo salar*, Sasa]; orange = Holostei [*Amia calva*, Amca and *Lepisosteus oculatus*, Leoc]). For holosteans, spotted gar sequences are indicated in brown and bowfin sequences are indicated in orange. The following sequence identifiers are abbreviated in the figure: Leoc LG29:293175 = Leoc LG29:293175-293459; Leoc JH591438 = Leoc JH591438:384055-384327; Leoc 00000000830 = Leoc ENSLOCG00000000830; Leoc LG29:151548 = Leoc LG29:151548-151823; Leoc 00000000745 = Leoc ENSLOCG00000000745; Leoc LG29:252003 = Leoc LG29:252003-252305; Leoc LG29:251128 = Leoc LG29:251128-251469; Leoc LG29:261282 = Leoc LG29:261282-261614; Lach JH130480.1 = Lach JH130480.1:43891-44139; Lach JH130632.1 = Lach JH130632.1:37775-37984; Ctid CI01000243:00274762 = Ctid CI01000243:00274762-00275673; Ctid CI01000243:00284609 = Ctid CI01000243:00284609-00301594; and Cyca LG38:21616820 = Cyca LG38:21616820-21618149. Scale bar indicates substitutions per million years.

### Bowfin DICP diversity is not the result of a lineage-specific expansion

All DICP Ig domains identified from the bowfin reference genome can be classified as either a D1 or D2 based on strong phylogenetic support [Bootstrap support (BSS) = 100]. Our phylogenetic analyses based on Ig domains from bowfin, aligned to DICP D1 and D2 domains from zebrafish, carp, salmon, pufferfish, tilapia, spotted gar and coelacanth, place the evolutionary history of holostean DICPs and the pairing of D1 and D2 domains into the broader context of vertebrate DICPs (**Fig. 4**). Within teleosts, the most detailed work on DICPs has been conducted within cyprinids (e.g., zebrafish, carp; Haire et al. 2012; Rodriguez-Nunez et al. 2016; Gao et al. 2018). It has been hypothesized that clusters of teleost DICP genes arose as a consequence of within-clade tandem gene duplication, largely through species-specific diversification (Haire et al. 2012), a result also supported by our phylogenetic analyses (**Fig. 4**). Holosteans depart from this pattern. Within each clade of holostean DICPs, intraspecific gene diversification events are less frequent than those observed in cyprinds. Instead, each DICP clade comprises a few gar and bowfin genes with no support for a monophyletic cluster of holostean genes (**Fig. 4**). Given that we have investigated the genomes of bowfin and spotted gar as well as a range of bowfin transcriptomes (see below), it is unlikely we have missed major clusters of DICPs. These results suggest that the DICP evolution in holosteans contrasts with that of teleosts such as zebrafish. Furthermore, gar and bowfin diverged several hundred million years ago (Near et al. 2012b; Dornburg et al. 2014; Hughes et al. 2018), a divergence on par with that of birds and crocodiles (Nesbitt 2003; Alfaro et al. 2009; Prum et al. 2015; Fabbri et al. 2017). This divergence, coupled with the observation that holosteans appear to have some of the slowest rates of molecular evolution among vertebrates (Braasch et al. 2016; Takezaki 2018) suggests that rather than a consequence of within-species diversification, DICP diversity in living holosteans is the result of maintaining a diversity of genes with ancient evolutionary origins.

### Bowfin DICP transcripts predict inhibitory, activating, secreted and functionally ambiguous protein structures

The 21 DICP sequences predicted from the bowfin reference genome include seven genes predicted to possess an extracellular D1-D2 organization and four genes predicted to possess an extracellular D1a-D2a-D1b-D2b organization (**Fig. 5**) with the remaining DICP genes reflecting partially annotated genes or pseudogenes (**Supplementary Table S1 and Fig. S1**). Two of the eleven predicted bowfin DICPs possess a cytoplasmic ITIM indicating a likely inhibitory function, whereas four DICPs possess a charged residue within their transmembrane reflective of an activating function, and five DICPs lack any identifiable signaling motif.

**Fig. 5.**
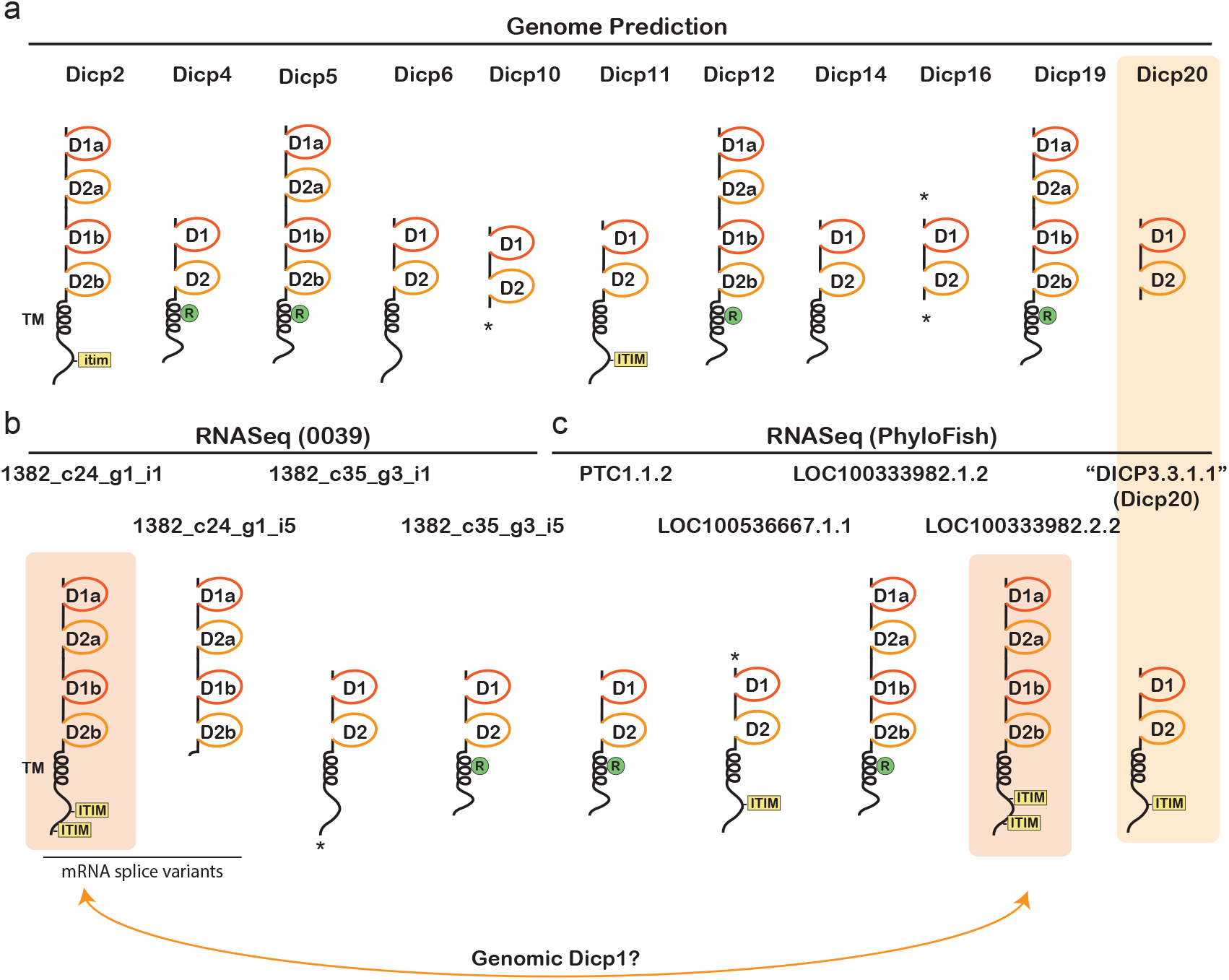
Predicted DICP protein architecture. Select DICP protein structures reflect sequences identified from **a** the bowfin reference genome, **b** RNA-seq from a single bowfin (0039) and **c** the PhyloFish database. Protein domains include: Ig D1 and D2 domains (orange), transmembrane domains (TM), cytoplasmic ITIM and ITIM-like (itim) sequences. The presence of a charged arginine within a TM indicates a potential activating receptor and is indicated by a green circle. Sequences that are predicted to be truncated at the 5’ and/or 3’ are indicated by asterisks (*). Shading of Dicp20 and “DICP3.3.1.1” indicate that they may reflect the same gene (see **Supplementary Note 2** and **Fig. S4**). Note that PhyloFish transcript names do not correspond to the genome-based DICP gene nomenclature. Shading of 1382_c24_g1_i1 (from bowfin 0039) and LOC100333982.2.2 (from PhyloFish) indicate that they likely reflect the same gene product and may represent Dicp1 (see **Supplementary Note 2** and **Fig. S9**).

In an effort to validate these predicted protein structures, BLAST searches of publicly available bowfin transcriptomes (**Supplementary Note 1**) were conducted and nine distinct transcripts identified (**Fig. 5 and Supplementary Table S2 and Fig. S2**). Specifically, tBLASTn searches identified five predicted bowfin DICP transcripts on the PhyloFish database (“DICP3.3.1.1”, LOC100536667.1.1, LOC100333982.1.2, LOC100333982.2.2, and PTC1.1.2) and revealed four distinct bowfin DICP transcripts from bowfin 0039 (1382_c24_g1_i1, 1382_c24_g1_i5, 1382_c35_g3_i1 and 1382_c35_g3_i5). All nine transcripts encode either a D1-D2 or D1a-D2a-D1b-D2b organization which is reflective of most cyprinid DICPs, although D1-only DICPs have been described from zebrafish and carp (Haire et al. 2012; Gao et al. 2018). As with teleost DICPs, bowfin DICPs include inhibitory, activating, secreted and functionally ambiguous forms.

### DICP sequence diversity indicates gene content variation

Gene content variation and alternative mRNA splicing appear to contribute to bowfin DICP diversity. None of the bowfin DICP transcripts mapped onto the reference genome with 100% accuracy. As the reference genome and transcriptome sequences were derived from different individuals, this observation indicates intraspecific DICP sequence variation in bowfin as described in zebrafish (Rodriguez-Nunez et al. 2016). Nevertheless, a phylogenetic comparison of transcript-encoded DICP D1 and D2 domains to genome-encoded D1 and D2 domains revealed multiple similarities (**Supplementary Fig. S3**). Direct comparisons of highly similar sequences suggest that certain DICP transcripts may reflect polymorphic versions of specific DICP genes and indicate that alternative mRNA splicing can also contribute to DICP diversity (summarized in **Fig. 5** with details provided in **Supplementary Note 2** and **Figs. S4-S9**).

### The number of bowfin NITR genes rivals that of teleosts

We previously reported fifteen distinct NITR sequences that occur within two genomic regions of the spotted gar genome (Braasch et al. 2016). We expanded this by identifying two additional gar NITR sequences (**Fig. 6a**) (Wcisel et al. 2017). Here we report a total of 34 bowfin NITR genes and pseudogenes (*nitr1* - *nitr34*) encoded across pseudochromosome Aca_scaf_8 and six unplaced scaffolds (**Supplementary Table S3**). The cluster on Aca_scaf_8 encodes the largest number of NITRs (*nitr1* - *nitr28*) in two clusters spanning approximately 313 Kbp and 42 Kbp (**Fig. 6a**). Each of the six smaller scaffolds that encode NITR sequences [Aca_scaf_68 (7420 bp), Aca_scaf_85 (5,185 bp), Aca_scaf_148 (3,624 bp), Aca_scaf_149 (3,603 bp), Aca_scaf_657 (1,711 bp), and Aca_scaf_1322 (1,237 bp)] encode one additional NITR. Teleost and gar NITRs genes typically encode several defining regions: a signal peptide sequence, a V-type Ig domain, an I-type Ig domain, J segments, a transmembrane domain with subsequent exons encoding a cytoplasmic tail (Yoder 2009). Our results demonstrate that many bowfin NITRs encode similar regions (**Fig. 6b** and **Supplementary Table S3)**.

**Fig. 6.**
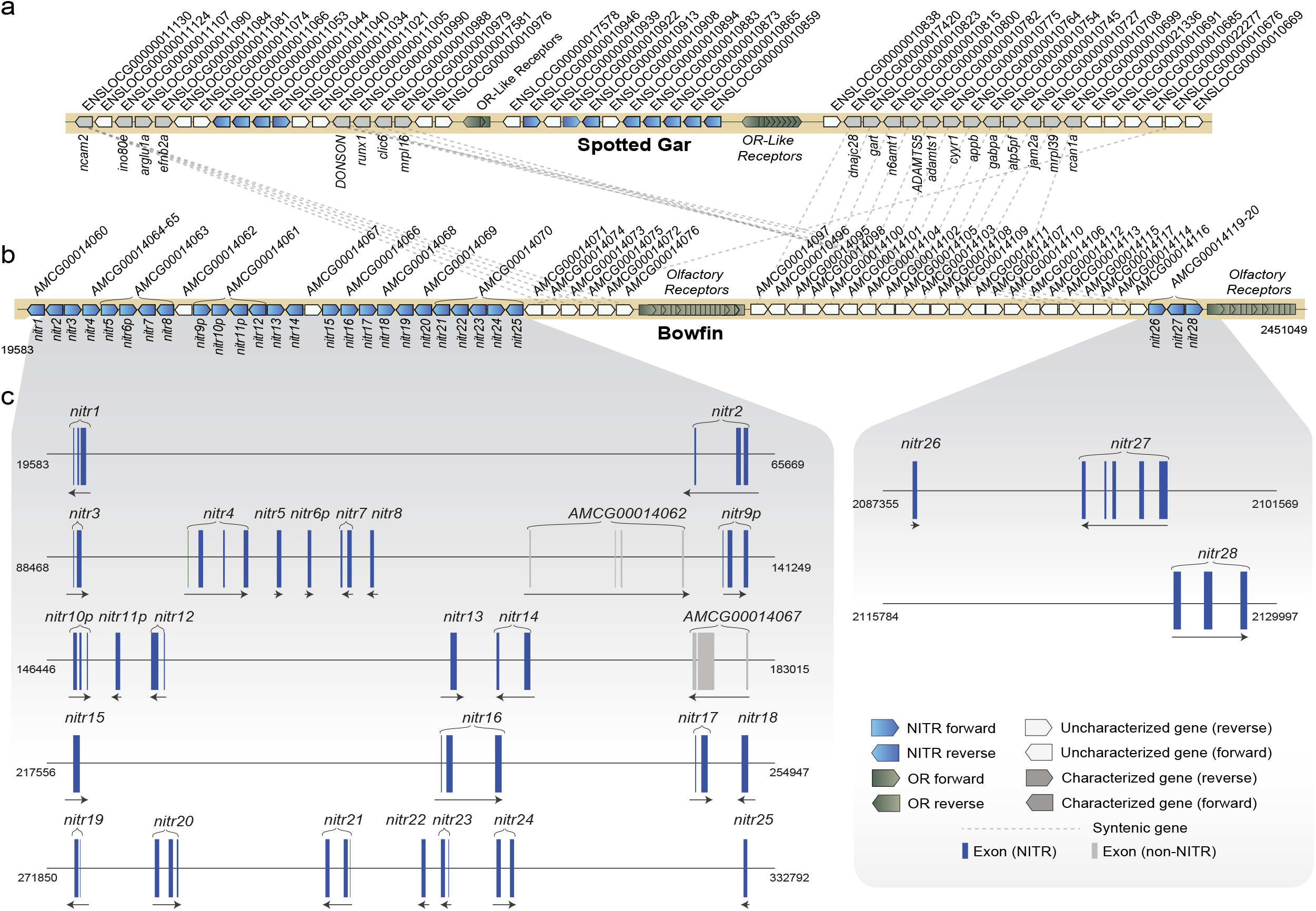
Bowfin NITRs are encoded on scaffold Aca_scaf_8. The syntenic relationship between the NITR gene clusters on spotted gar linkage group 14 **(a)** and bowfin pseudochromosome Aca_scaf_8 **(b)** is indicated. A number of genes flanking the NITR clusters (e.g. *runx1, donson*, and a number of olfactory receptors) are conserved in both holostean species. Each pentagon reflects a single gene with transcriptional orientation indicated. Gene sequence identifiers are shown above the genes (ENSEMBL for gar and AMCG for bowfin) with common gene symbols below. **(c)** A detailed exon map represents the current knowledge of NITR genomic organization within the reference genome. Nucleotide position within Aca_scaf_8 is indicated on the left and right of each genomic region.

Although not all bowfin NITR genes can be confirmed to encode functional proteins (e.g. pseudogenes *nitr9p and nitr30p* encode an internal stop codon or frameshift within the V domain and *nitr10p* and *nitr11p* encode partial I domains), at least six genes (*nitr2, nitr16, nitr20, nitr21, nitr24*, and *nitr34*) are predicted to encode both a V and I domain and are considered functional NITRs (see below and **Supplementary Table S3** and **Fig. S10**). Because *nitr27* encodes a V domain with only part of an I domain (it is missing the C-terminal region of the I domain including the highly conserved C^104^ that stabilizes the Ig fold), we exclude it from phylogenetic analyses of I domains. Due to the large number of ambiguous base (N) assignments in the assembly, several partial NITR genes (encoding 1 or 2 exons) were identified in Aca_scaf_8 that may reflect pseudogenes or partial genes (e.g. *nitr3, nitr7, nitr8, nitr13*, etc.) Nevertheless, the identification of 34 NITR sequences in bowfin is on par with the 39, 44 and 30 NITR genes identified in zebrafish, medaka and sea bass, respectively (Yoder et al. 2004; Desai et al. 2008; Ferraresso et al. 2009; Rodriguez-Nunez et al. 2016) suggesting that the large number of teleost NITRs is not just a result of paralog retention following the TGD.

### NITRs are likely derived from an ancient gene family that gave rise to V(D)J recombination

Teleost and gar NITR V domains are highly similar to TCR and Ig gene V domains. However, in both lineages NITR I domains possess six highly conserved cysteines (two cysteines that form the disulfide bond promoting the Ig-fold and four novel cysteines that have not been identified in any other class of Ig domain-containing protein family) that provide a means to distinguish NITRs from TCR and Ig genes (Yoder 2009; Wcisel and Yoder 2016) (**Fig. 7)**. We demonstrate that bowfin NITR Ig domains possess an architecture similar to other NITR V and I domains. Alignment-based comparisons between bowfin, gar, and teleost V domains (**Fig. 8)** reveals the presence of residues that are conserved across NITR, T-cell receptor (TCR) and Immunoglobulin Ig domains such as C^23^ and C^104^ (Litman et al. 2001; Yoder 2009). In comparison to the V domains, the alignment of the bowfin I domains reveals striking within-species sequence conservation (**Fig. 7)** We also find that about half of bowfin NITR V domains and the majority of the NITR I domains possess nearly perfect germ-line joined consensus J sequences, FGxGTxLx(V/L).

**Fig. 7.**
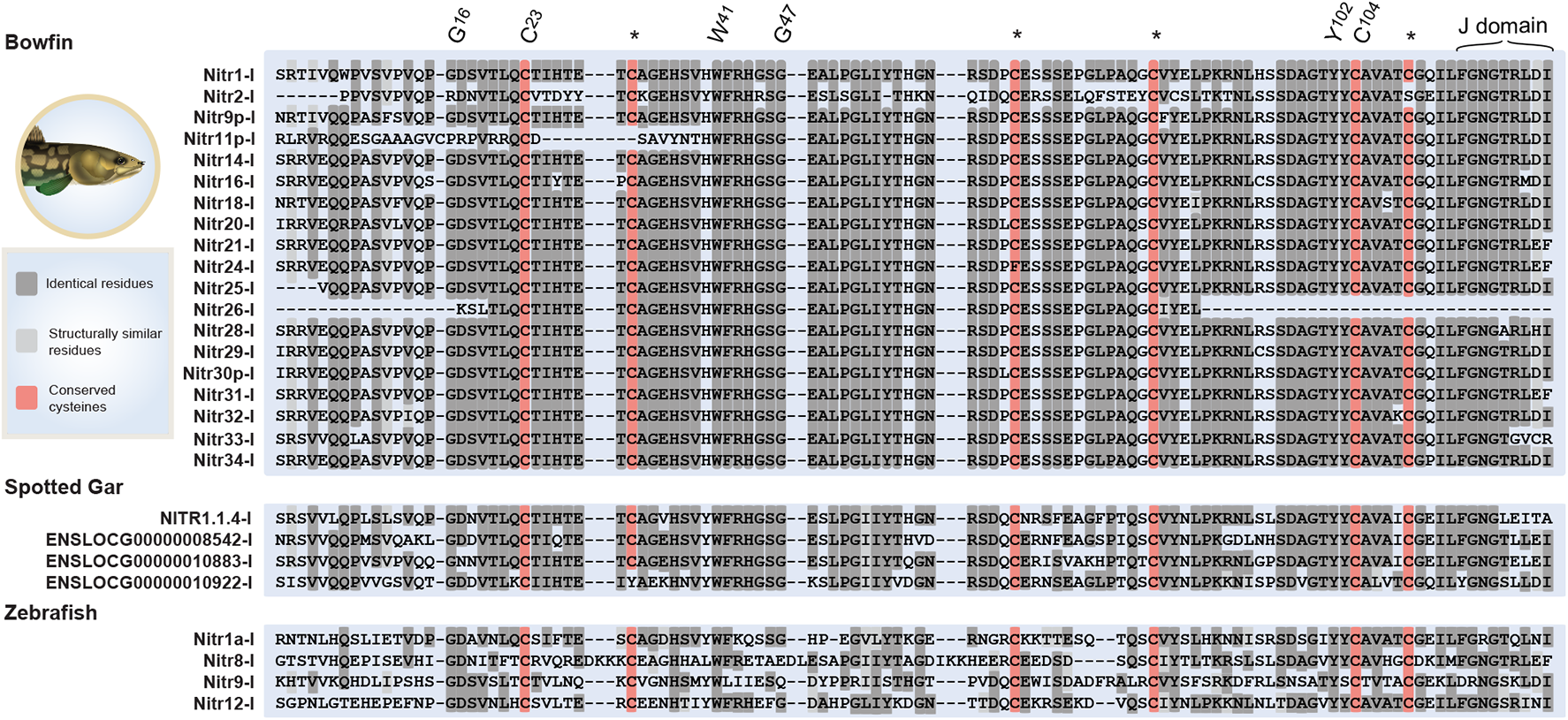
Conservation of the NITR I domain in bowfin demonstrates I-J motifs in both species of holosteans. Protein sequence alignment of the I region of bowfin NITRs compared with representative spotted gar and zebrafish sequences with the conserved J domain indicated for all species. Sequences are shaded by similarity with dashes indicating gaps in the alignment. Identical residues are shaded dark gray; structurally similar residues shaded light gray; and conserved cysteines shaded red. Key residues are numbered using the IMGT numbering system (Lefranc et al. 2015). Asterisks (*) indicate cysteines indicative of a NITR I domain.

**Fig. 8.**
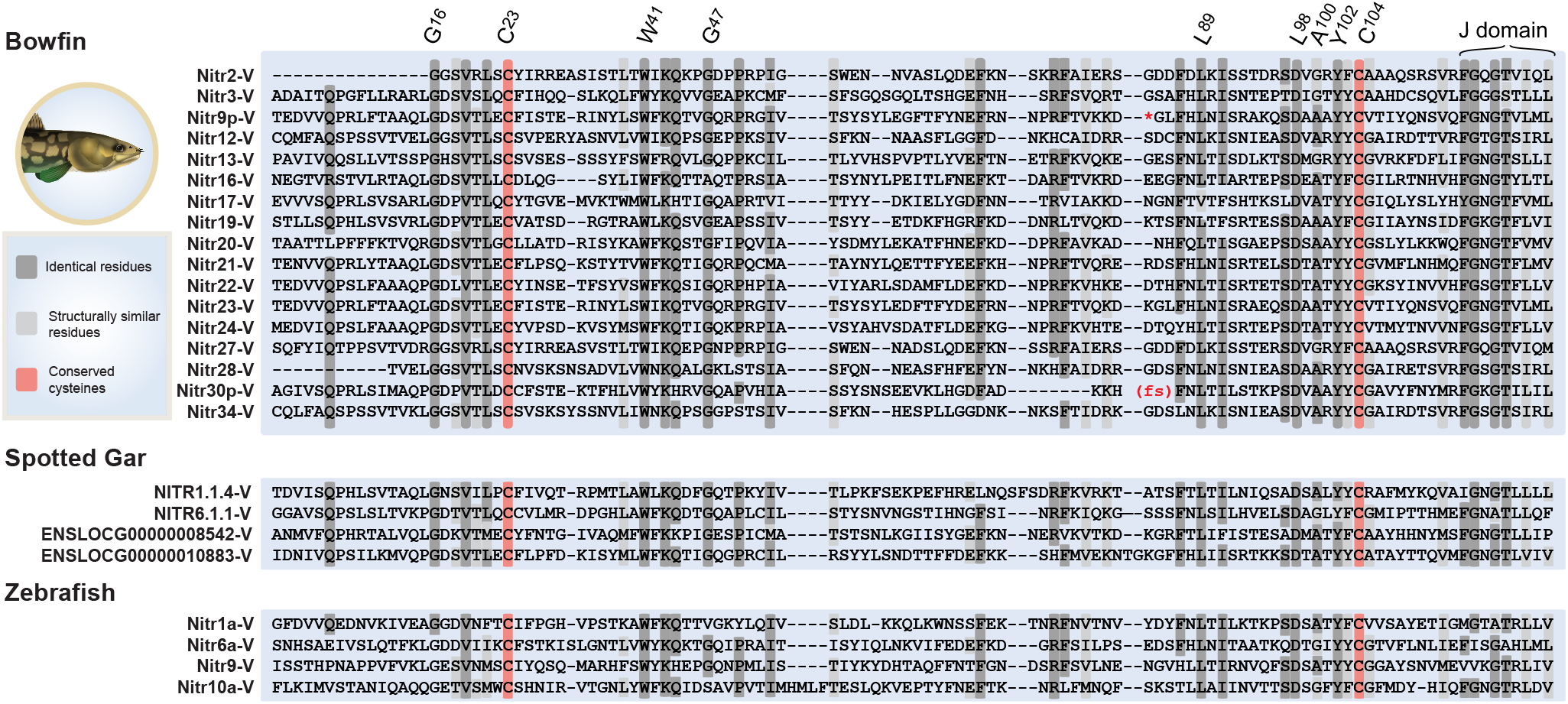
Conservation of the NITR V domain in bowfin demonstrates V-J motifs in both species of holosteans. Protein sequence alignment of the V region of bowfin NITRs compared with representative spotted gar and zebrafish sequences with the conserved J domain indicated for all species. Sequences are shaded by similarity with dashes indicating gaps in the alignment. Identical residues are shaded dark gray; structurally similar residues shaded light gray; and conserved cysteines shaded red. An internal stop codon in Nitr9p-V is represented by an asterisk and a frame shift in Nitr30-V is represented by (fs). Key residues are numbered using the IMGT numbering system (Lefranc et al. 2015).

The presence of J or J-like sequences in a single exon with a V or I domain is characteristic of teleost NITRs, and supports the proposal that NITRs either represent an ancient gene family that may have given rise to V(D)J recombination in the adaptive immune system, or an evolutionary novelty that arose as a consequence of the teleost genome duplication (Strong et al. 1999; Litman et al. 2001, 2003; Yoder et al. 2004). The hypothesis that NITRs arose as a consequence of the TGD was supported by the lack of evidence for NITR homologs in cartilaginous fishes (Yoder et al. 2004). However, our finding of V-J and I-J motifs in both species of holosteans rejects this TGD-derived hypothesis. Instead, NITRs are in fact an ancient family of innate immune receptors with origins that predate the TGD. Future investigations focused on the origins of NITRs are needed to determine if this gene family is unique to neopterygians, or represents an older gene family that has been maintained in ray-finned fishes but subsequently lost in sarcopterygians.

### Holostean NITR diversity is derived from lineage specific expansions

Although NITR’s may represent an ancient gene family, NITR sequence diversity appears largely species-specific, meaning any individual NITR sequence is generally more similar to another NITR sequence from within the same species than another (Desai et al. 2008; Yoder 2009; Ferraresso et al. 2009). Our phylogenetic analyses place bowfin NITR diversity within the broader context of neopterygian sequences and support this hypothesis. Analysis of the I domain reveals that the vast majority of bowfin NITRs are monophyletic. The sole exception is Nitr2, which is resolved as the sister lineage (BSS = 91) to a well-supported clade (BSS = 97) that contains all spotted gar NITRs. In both holosteans and teleosts, within-species receptor diversity largely evolved *in situ* **(Fig. 9)**. However, the *in situ* diversification of gar and bowfin NITR I domains represents a dramatic shift towards estimated branch lengths smaller than those estimated in teleosts. This contrast is of particular note as the estimated time of divergence between gar and bowfin is either similar to or slightly exceeds the time to common ancestry for the teleosts in this study (Near et al. 2012b; Dornburg et al. 2014; Hughes et al. 2018). This contrast between holostean and teleost branch lengths is also evident in a phylogenetic analysis of V domains (**Fig. 10**). Given the high rate of sequence evolution in NITR V domains versus I domains, our phylogenetic analyses reveal numerous cases of likely evolutionary convergences in V-domain sequence similarity between teleosts as well as teleosts and holosteans (**Fig. 10**). These results raise the question of whether the teleost genome duplication catalyzed faster rates of molecular evolution in teleosts, or if other mechanisms underlie the slow rates of sequence evolution that characterize holosteans (Braasch et al. 2016). We hope to address this question in the future with a broader sampling of species and innate immune receptors.

**Fig. 9.**
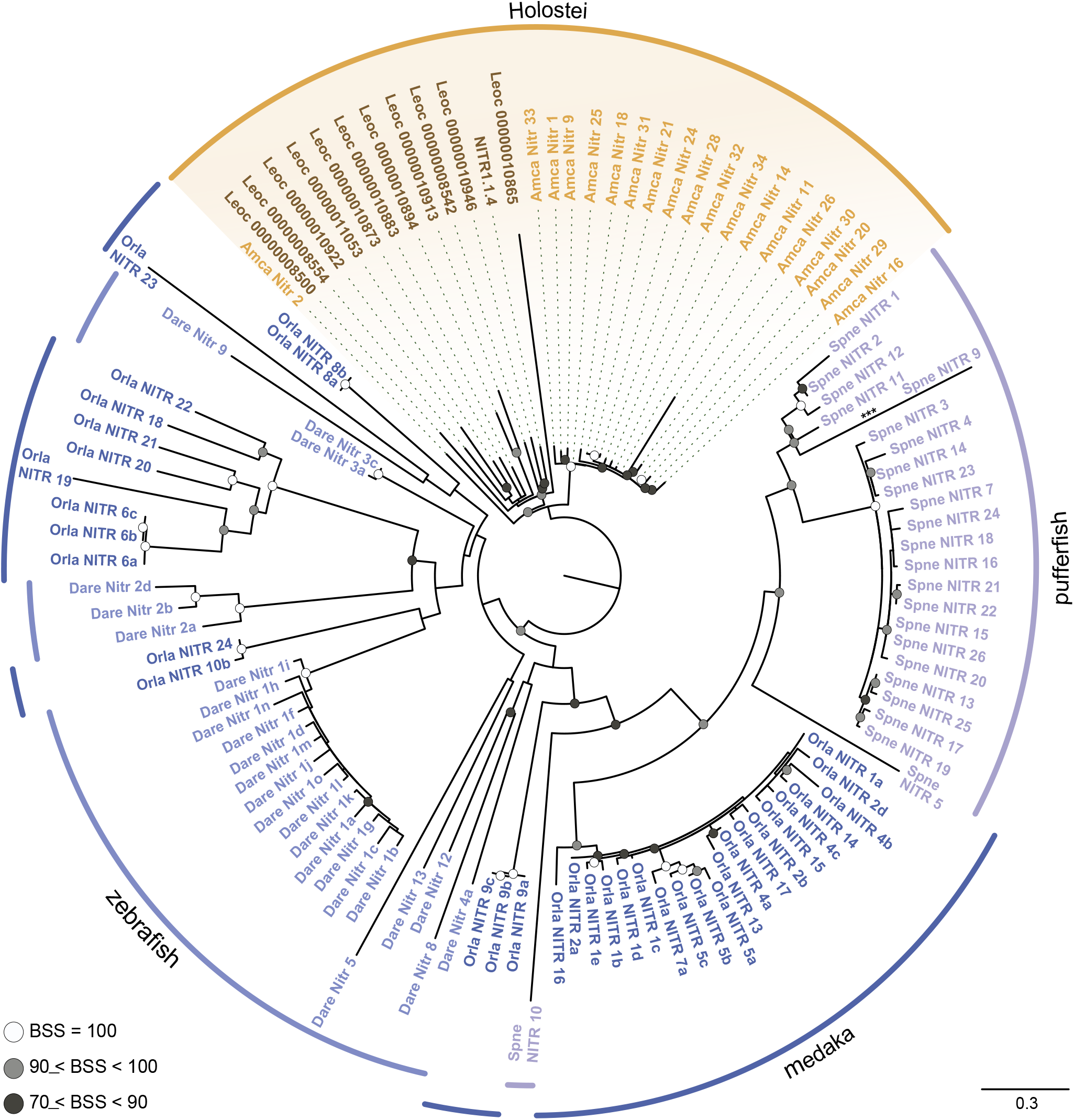
NITR I domain diversification mirrors lineage-specific gene expansions observed in teleosts, but with slower molecular rates. Maximum likelihood phylogeny of NITR I domains inferred using IQ-TREE. Circles at nodes indicate bootstrap support values (BSS) with filled black circles black indicating BSS=100, gray circles indicating BSS values greater than 90 but less than 100, and white circles indicating BSS values greater than 70 but less than 90. Lineages are indicated by the color coded arcs surrounding the tree and colored taxon labels (light blue = zebrafish [*Danio rerio*, Dare]; dark blue = medaka [*Oryzias latipes*, Orla]; purple = pufferfish [*Sphoeroides nephelus*, Spne]; orange = Holostei [*Amia calva*, Amca and *Lepisosteus oculatus*, Leoc]:). For holosteans, spotted gar sequences are indicated in brown and bowfin sequences are indicated in orange. Gar ENSEMBL sequence identifiers are abbreviated by removing “ENSLOCG” from the identifier in the figure (e.g. “Leoc ENSLOCG00000008554” is abbreviated “Leoc 00000008554”). Scale bar indicates substitutions per million years and asterisks (***) indicates branch length scaled by 50% for visualization.

**Fig. 10.**
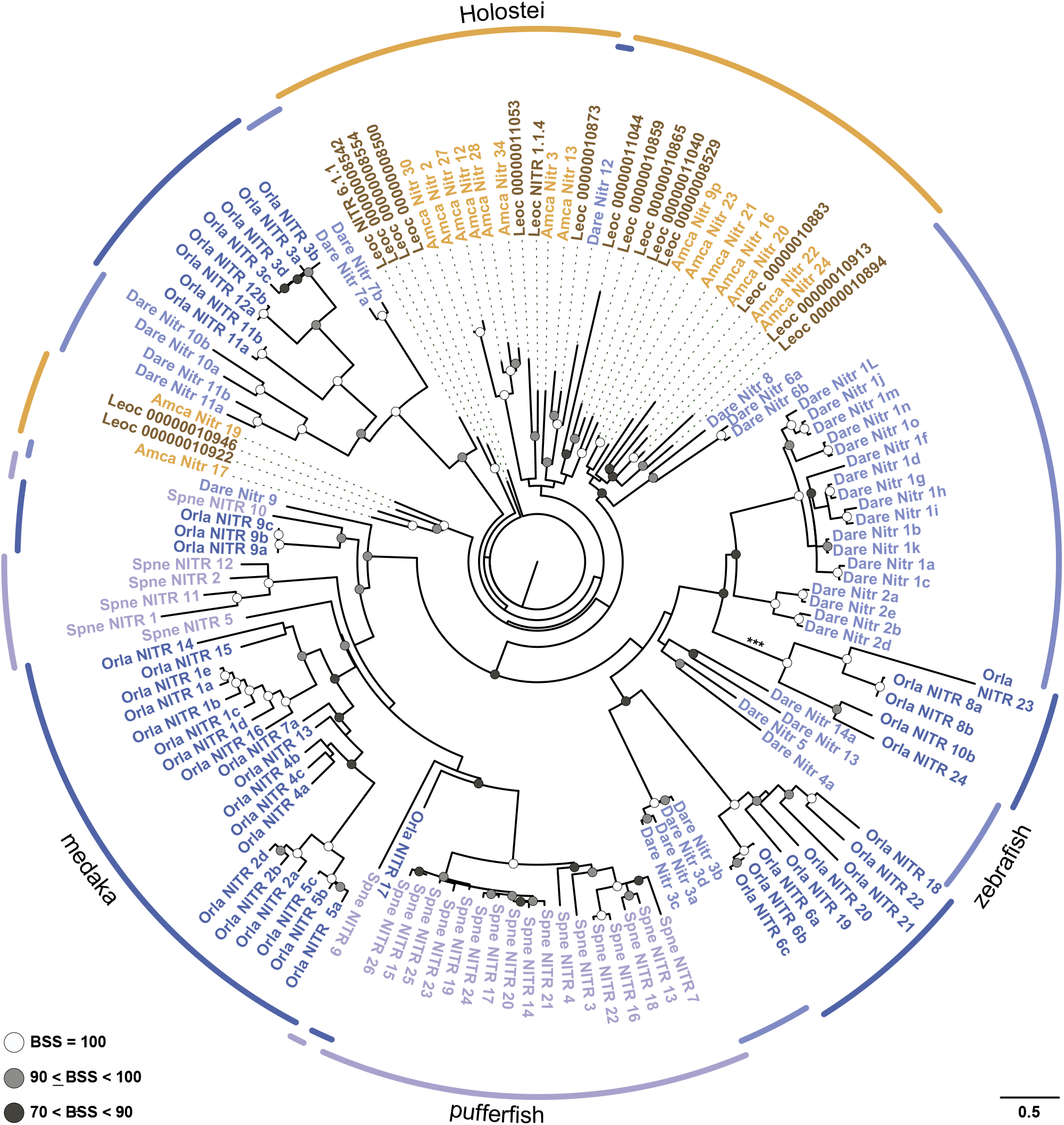
Holostean NITR V domain diversification is slower than that of teleosts. Maximum likelihood phylogeny of NITR V domains inferred using IQ-TREE. Circles at nodes indicate bootstrap support values (BSS) with filled black circles black indicating BSS=100, gray circles indicating BSS values greater than 90 but less than 100, and white circles indicating BSS values greater than 70 but less than 90. Lineages are indicated by the color coded arcs surrounding the tree and colored taxon labels (light blue = zebrafish [*Danio rerio*, Dare]; dark blue = medaka [*Oryzias latipes*, Orla]; purple = pufferfish [*Sphoeroides nephelus*, Spne]; orange = Holostei [*Amia calva*, Amca and *Lepisosteus oculatus*, Leoc]). For holosteans, spotted gar sequences are indicated in brown and bowfin sequences are indicated in orange. Gar ENSEMBL sequence identifiers are abbreviated by removing “ENSLOCG” from the identifier in the figure (e.g. “Leoc ENSLOCG00000008554” is abbreviated “Leoc 00000008554”. Scale bar indicates substitutions per million years and asterisks (***) indicates branch length scaled by 50% for visualization.

### Predicted functional diversity in bowfin NITRs matches that of teleosts

Of the 34 NITR sequences we manually predicted and annotated from the bowfin reference genome six are predicted to possess an extracellular V-I organization (**Fig. 11a** and **Supplementary Fig S10**.) with the remaining NITR genes reflecting partially annotated genes or pseudogenes (**Supplementary Table S3**). The majority of the sequences identified by automated prediction software do not reflect NITR protein architecture described in teleosts and are not supported by transcriptome data (below). The one exception is AMCP00014119 that is predicted to encode a bonafide NITR (Nitr27), albeit with a truncated I domain.

**Fig. 11.**
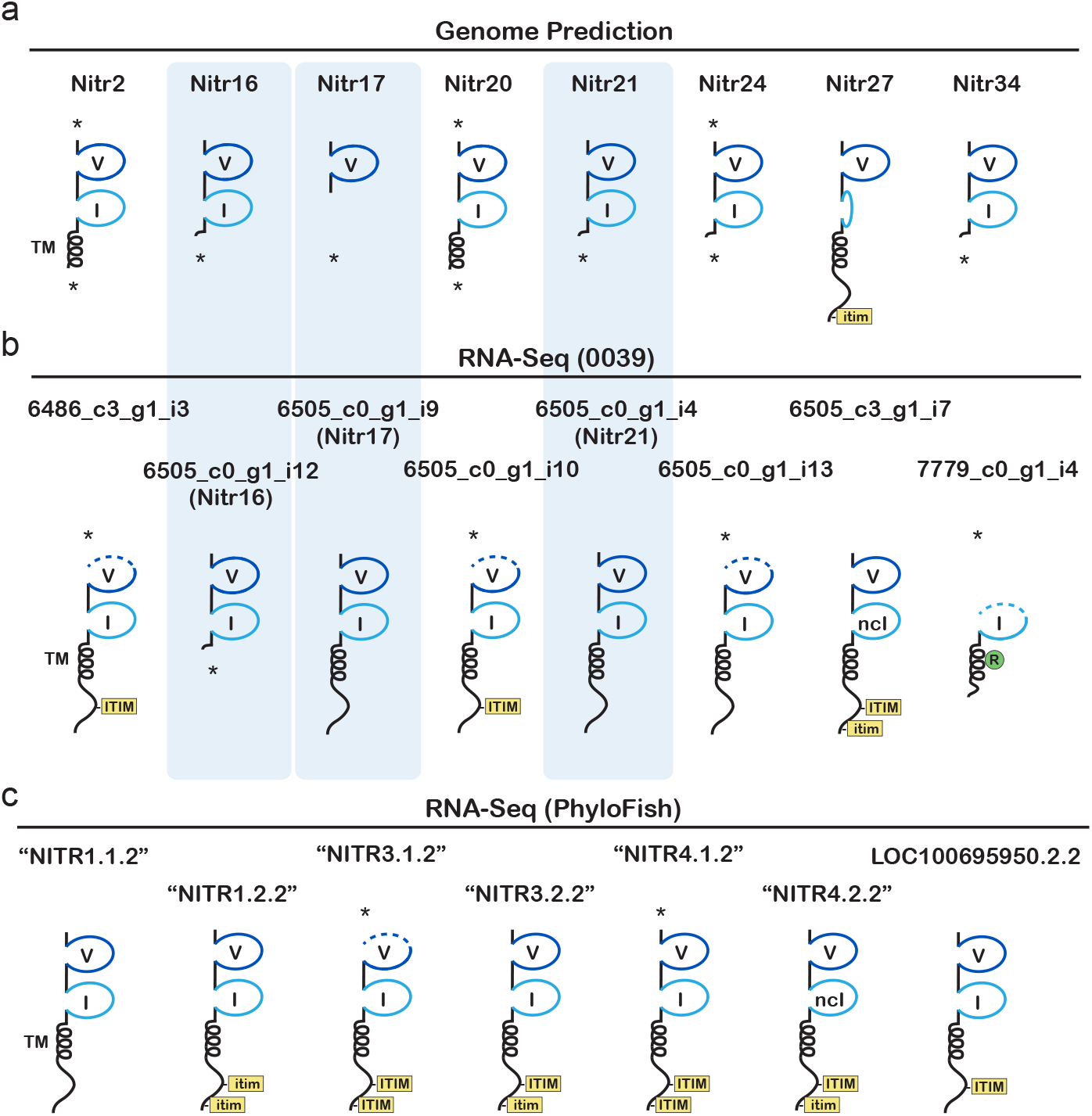
Predicted NITR protein architecture. Select NITR protein structures reflect sequences identified from **a** the bowfin reference genome and from **b** RNA-seq from a single bowfin (0039) and **c** from the PhyloFish database. Protein domains include: Ig V and I domains (blue), transmembrane domains (TM), cytoplasmic ITIM and ITIM-like (itim) sequences. The presence of a charged arginine within a TM indicates a potential activating receptor and is indicated by a green circle. Sequences that are expected to be truncated at the 5’ and/or 3’ are indicated by asterisks (*). Nitr27 encodes a truncated I domain which is shown as a compressed Ig domain. Shading of Nitr16 with 6505_c0_g1_i12, Nitr17 with 6505_c0_g1_i9, and Nitr21 with 6505_c0_g1_i4 indicate that they may reflect the same gene (see **Supplementary Note 3** and **Figs. S13-S15**). Note that PhyloFish transcript names do not correspond to the genome-based NITR gene nomenclature.

As described for DICPs, we searched for bowfin NITR transcripts with BLAST searches of publicly available bowfin transcriptomes (**Supplementary Note 1**) and 26 distinct transcripts were identified (**Fig. 11b-c** and **Supplementary Table S4 and Fig. S11**). Fourteen of these transcripts encode both a V and an I domain which reflects what is predominantly observed in teleosts (Litman et al. 2001; Yoder 2009; Ferraresso et al. 2009). One transcript, 7779_c0_g1_i4 is truncated but predicted to encode an activating NITR by the presence of a charged residue within the transmembrane domain (**Fig. 11b and Supplementary Fig. S11**). The majority of the other bowfin NITR transcripts with both V and I domains encode ITIM- or itim-like sequences and are considered inhibitory receptors. As V-only NITR genes have been described in some teleosts (Yoder et al. 2004; Desai et al. 2008; Ferraresso et al. 2009), and an alternatively spliced I-only NITR has been described in zebrafish (Shah et al. 2012), we report similar bowfin transcripts in **Supplementary Table S4** and **Fig. S11**. As with teleost NITRs, bowfin NITRs appear to include inhibitory, activating, secreted and ambiguous forms.

### NITR sequence diversity indicates gene content variation

Gene content variation appears to contribute to bowfin NITR diversity. As with DICPs, none of the bowfin NITR transcripts mapped onto the reference genome with 100% accuracy. A phylogenetic comparison of transcript-encoded NITR V domains to genome-encoded V domains revealed multiple similarities (**Supplementary Fig. S12**), however, only eight of 26 bowfin transcripts could be mapped back onto the reference genome with some level of certainty (**Supplementary Note 3, Table S4** and **Figs. S11, S13-S15)**. As described in **Supplementary Note 1**, the reference genome and transcriptome sequences were derived from different individuals indicating that bowfin NITRs, like zebrafish NITRs and bowfin DICPs (see above), display intraspecific gene content variation. More specifically, all eight of these transcripts were derived from a single bowfin (0039) that was collected near the geographic location where the individual used for the reference genome was collected (**Supplementary Note 1**) (Thompson et al. 2021).

## Summary: Considering DICPs and NITRs in the context of the TGD

Over the past several decades, numerous ray-finned fish genes and gene families have been hypothesized to be teleost-specific due to the lack of reference genomes for the few living species of non-teleost ray-finned fishes and the absence of these genes in more distantly related sarcopterygians (lungfish, coelacanth, tetrapods) or earlier diverging non-bony vertebrates (lamprey, hagfish, chondrichthyans). However, the continued sequencing of non-teleost ray-finned fishes provides us with an exciting opportunity to test expectations of TGD-catalyzed innovation and contextualize general principles of genome evolution. Our results illuminate the functional diversity of NITR and DICP receptors in bowfin **(Figs. 5** and **11)**, suggesting that high functional diversity of these receptors represent a hallmark of neopterygians if not actinopterygians more generally. These findings do not support the hypothesis that the TGD catalyzed a sudden burst of evolutionary novelty that gave rise to new teleost fish innate immune receptor families. Instead, this diversity was already in place and likely has far more ancient origins within ray-finned fishes.

Combining our results with other investigations of the bowfin and gar genomes (Braasch et al. 2016; Wcisel et al. 2017; Thompson et al. 2021), it is evident that rather than catalyze a pulse of molecular diversification, the TGD likely had a dramatic impact on the overall architecture of the teleost genome that may have provided the substrate for subsequent within-lineage diversification. We find that the DICP genes are located within two closely positioned clusters on the same pseudochromosome (Aca_scaf_14) as gene clusters of MHC class I lineages (U, Z, P, and L) as well as MHC class II and class III genes, even though some MHC I class I genes (P, L, S and H lineages) are present on other pseudochromosomes. This condition contrasts with that of teleost fishes such as zebrafish where a cluster of DICP genes is linked to a cluster of MHC class I Z lineage genes, and U lineage genes are on other chromosomes (Dirscherl et al. 2014; Rodriguez-Nunez et al. 2016). Our results suggest that the fragmentation of MHC class I/II/III genes and DICP genes in teleosts arose due to differential loss of genes from paralogous chromosomes in combination with post-TGD chromosomal rearrangements. Additional evidence for this hypothesis is found in Aca_scaf_8: here bowfin NITR genes are intermingled with other Ig-domain containing proteins such as CD276-like (AMCP00014071) and NCAM2-like sequences (AMCP00014072, AMCP00014074, AMCP00014075, AMCP00014076) (**Fig. 6A**). Conserved gene synteny between spotted gar linkage group 14 and bowfin Aca_scaf_8 is observed, exemplified by the presence of a cluster of olfactory receptor (OR) genes as well as a number of single copy genes (e.g. *runx1, DONSON*, etc). This synteny is noteworthy, as the NITR region of the spotted gar genome was not found to be syntenic with model teleosts such as zebrafish thereby suggesting a loss of synteny between the NITR loci of teleosts and holosteans following the TGD.

Collectively, our results align with an emerging perspective concerning the fate of paralogs in genome evolution that predicts paralog diversity to rarely be maintained over deep stretches of evolutionary time (Nei and Rooney 2005; Ferraresso et al. 2009; Inoue et al. 2015; Fernández and Gabaldón 2020). There is little reason to expect that a sudden shift in the diversification of a gene family should occur in the absence of an external catalyst that imposes strong selective pressures. Instead it is far more likely that the clustered organization of these and similar families of innate immune receptors have, and continue to provide the genomic substrate required to persist in the face of evolving pathogenic threats over several hundred million years of teleost evolution. The changes in pathogen driven selective pressures as ray-finned fishes have made evolutionary transitions to novel biomes such as saltwater to freshwater transitions (Yamanoue et al. 2011; Nakatani et al. 2011; Davis et al. 2012), invaded new adaptive zones (Dornburg et al. 2011; Burns and Sidlauskas 2019; Friedman et al. 2019), or faced changes in climatic conditions (Near et al. 2012a; Siqueira et al. 2019), could be associated with pathogenic spread. Transitions such as these may explain the high within-lineage diversity of these receptor families. In particular, it is possible that these clusters are hot spots for gene birth and death, which could provide the mechanism for our observations of intra-specific gene content variation. Although the diversification dynamics of these gene clusters remain unknown, inter-individual variation in gene content would provide a wider degree of protection to the next unknown pathogen (Uhrberg et al. 2002; Vilches and Parham 2002; Tukwasibwe et al. 2020) that at the level of meta-populations could mitigate the impact of novel pathogens during such evolutionary events. As we expand our ability to move detailed comparative genomic studies from the root of the teleost phylogeny to the tips, testing how such shifts in ecological opportunities often associated with the diversification of lineages (Near et al. 2013; Berv and Field 2018) and key ecological phenotypes (Salzburger 2018; Daane et al. 2019) has shaped the genomic basis of ray-finned fish immunity represents an exciting research frontier.

## Acknowledgments

We thank Thomas Near (Yale University) for helpful discussions about bowfin biology and the consequences of the teleost genome duplication and Madhusudhan Gundappa (@fish_lines) for illustrations of bowfin and spotted gar.

## Authors’ contributions

AD and JAY conceived of and designed the study. EF, KZ, AWT, IB, LRA, TO, DJW, and AWT assembled sequence data. AD, JAY, TO, EF, LRA, KZ, AWT, IB, and DJW analyzed the data. AD and JAY wrote the first draft of the manuscript. All authors contributed to the writing of the final manuscript.

## Data availability

All data used in this study are publicly available on NCBI, Ensembl, or the PhyloFish database (http://phylofish.sigenae.org/). Publicly available bowfin sequences accessed during this study are also provided as full length fasta formatted sequences and partitioned sequences linked to the delimitations of domains (e.g., NITR-I, NITR-V, etc) in the Supplementary Materials.

