## Supplementary Notes and Figures for "Holosteans contextualize the role of the teleost genome duplication in promoting the rise of evolutionary novelties in the ray-finned fish innate immune system"

a. Department of Bioinformatics and Genomics, University of North Carolina at Charlotte, Charlotte, NC USA
b. Department of Molecular Biomedical Sciences, North Carolina State University, Raleigh, NC, USA
c. North Carolina Museum of Natural Sciences, Raleigh, NC, USA
d. Department of Integrative Biology and Ecology, Evolution, and Behavior Program, Michigan State University, East Lansing, MI, USA
e. Department of Evolutionary Studies of Biosystems, SOKENDAI (Graduate University for Advanced Studies), Hayama, Japan
f. Comparative Medicine Institute, North Carolina State University, Raleigh, NC, USA
g. Center for Human Health and the Environment, North Carolina State University, Raleigh, NC, USA

* **Corresponding authors at**:

**Supplementary Notes.**

*1. Bowfin transcriptome sources.*

Transcriptome databases used in this study include the PhyloFish database (Braasch et al, 2016; Pasquier et al. 2016) and RNA-Seq from an individual bowfin “0039” (Thompson et al 2021). The PhyloFish bowfin transcriptome database includes sequences from brain, liver, gills, heart, muscle, kidney, bones, intestine from one adult female; ovary from one adult female; and testes from one adult male (NCBI SRA accession number: SRP044783). The “0039” bowfin transcriptome database includes sequences from immune tissues (spleen, liver, gills and gut) of a single adult fish (NCBI SRA accession number SRR11303972, and TSA accession number GIOP00000000). All bowfin used in this study were collected from the same general area near Thibodeaux, Louisiana. The individual used for the reference genome was collected from 29.812569 N 91.220803 W and bowfin “0039” was collected from 29.929972 N 90.747389 W (Braasch et al, 2016; Thompson et al. 2021).

*2. DICP sequence variation.*

A phylogenetic comparison of transcript-encoded DICP D1 and D2 domains to genome-encoded D1 and D2 domains revealed multiple similarities (**Supplementary Fig. S3**). For example, the protein encoded by transcript “DICP3.3.1.1” overlaps Dicp20 with 100% identity leading to the prediction that this transcript is a product of this gene (note that “DICP3.3.1.1” is the PhyloFish transcript nomenclature) (**Supplementary Fig. S4**). Although transcripts 1382_c24_g1_i1 (identified from bowfin 0039) and LOC100333982.2.2 (identified from PhyloFish) encode nearly identical proteins, they also share high similarity to the product encoded by the partial gene product Dicp1 (**Supplementary Fig. S5**). Similarly the products of transcripts LOC100333982.1.2 and LOC100536667.1.1 share sequence homology with Dicp12 and Dicp14, respectively, but are different enough to question if they reflect the same genes (**Supplementary Figs. S6-S7**). In addition, the proteins encoded by transcripts 1382_c35_g3_i1 and 1382_c35_G3_i5, share identical signal peptide, D1 and D2 sequences but encode distinctly different transmembrane and cytoplasmic tails suggestive of alternative splicing, exon swapping or a recent gene duplication event. In addition, the partial sequences identified for Dicp9 and Dicp10 share nearly identical D1 and D2 domains with these transcripts (**Supplementary Fig. S8**). Finally, transcripts 1382_c24_g1_i1 and 1382_c24_g1_i5 are predicted to reflect alternative mRNA splice variants that share four extracellular Ig domains (D1a-D2a-D1b-D2b) with one encoding a secreted form (1382_c24g_g1_i5) and the other encoding a membrane bound form that includes cytoplasmic ITIMs (1382_c24_g1_i1) (**Fig 5** and **Supplementary Fig. S9**).

*3.* NITR sequence variation*.*

A phylogenetic comparison of transcript-encoded NITR V domains to genome-encoded V domains revealed multiple similarities (**Supplementary Fig. S12**), however, only eight of 26 bowfin transcripts could be mapped back onto the reference genome with some level of certainty (**Supplemental Table S4** and **Fig. S11)**. For example, the proteins encoded by transcripts 6505_c0_g1_i12, 6505_c0_g1_i9 and 6505_c0_g1_i4 are nearly identical to the partial protein sequences predicted from Nitr16, Nitr17 and Nitr21, respectively, indicating that these transcripts may reflect polymorphic variants of these genes (**Fig. 11a-b** and **Supplementary Figures S13-S15**).

*4. References.*

Braasch I, Gehrke AR, Smith JJ, et al (2016) The spotted gar genome illuminates vertebrate evolution and facilitates human-teleost comparisons. Nat Genet 48:427–437

Pasquier J, Cabau C, Nguyen T, et al (2016) Gene evolution and gene expression after whole genome duplication in fish: the PhyloFish database. BMC Genomics 17:368

Thompson A, Hawkins M, Parey E, et al (2021) The genome of the bowfin (Amia calva) illuminates the developmental evolution of ray-finned fishes. Nature Genetics (accepted). https://doi.org/10.21203/rs.3.rs-92055/v1

**>Dicp1 (partial)**

MRSAIFPLLISLSCLSEWAQRLGVEXXXXXXXXXXXXXXXXXXXXNSSLHIDRLRTRDTGQYTCHQYVNGKHYTSGLPVTLFLLSISVSSSSSACLRAGDRL
TLKCGLDCGGGAGSCSETPQGLTLSWRDESGVPPKDERDRYSITELRGVHSQLSVTLRQSDHNKSWTCVLTERGEMKTSESYTTTLSDELFSTVGRLLLLSC
VAPVSLGPGETLRWTYRQSRTSQAVVLYSLRSLTETPLKGTVPHSGRAVMSANSSLLIHNVQTGDAGLYCCQRDKDSGPKKTHRTFALNTLSXXXXXXXXXX
XXXXXXXXXXQLVLQPQSSERIRCSVERQGLERVSQDLTIKIEAGPGGKFPMVEVVSAAAAFCVLLLLVFIIFLVMKMKKTKKEGKKDRVPSDQAEERPESV
YHSPEEIMSCAAPPHTDD**VNYTSV**MFKSKKRGPEREQTFLPNSDD**VIYSAV**TTTQ

**>Dicp2 (AMCP00006279, modified)**

MACQSLLLFLFVPVWSTDSGLGQGSELVLGIKSSGDTLYSSVGGSAILPCVGVTHQHCPSTTWMLSRKFRPDTEVVKLGKVTDVERAGRLSVGSDCSLHIER
LKSPDAGVYTCSHRNTPGTSVTLHLLNITVSLQTDLKSGTEIELHCFISTSSAIEPCNPPPKGVRVGWVNGRGDALQGSRYTVRNESVCHSVLTLTLQPSDH
NTQWRCDLTQEGSQRVTSTFTTELTDGIQQVYRVVGSSVTLPCAINVTQTLGQGEKLEWFLRTWDDPIGHTLFSLSSDWKVSPINPTVVPGVKMAANSSLVL
PALSAQHSGIYLCSLYSGSSVKKELWRLALHIVEVSSDPTGGIQKGSGINLTCSLTCEDVCDKNTELTWRDSAGCSLQGGPAERVNNTLSSWLLVPELQSSE
RIWCSVVREGLERVEQDITVVVSAGGQAGLAVTAAVSSVAIIILLVCVVAIRTYMRKSRKGRSGAGQPADGVADLYEDCGDTSYAQVLFNGDRQQAERRRER
EDT**AAVYSL**LSAVN

**>Dicp4**

MRSPLFPLLISLSCLSEWAQRLGVEDELFSTVGRLLLLSCVAPVSLGPQETLRWTYRQSRTSQAVVLYSLRSLTETPLKGTVPHSGRAVMSANSSLLIHNVQ
TGDAGLYCCQRDKDSGPKKTHRTFALNTLSVSSNLSEEAQNGSAVTLTCSLTCGFDCEENTELIWRDSTGNSLQGGTSERNKSTISSQLVLQPQSSERIRCS
VEREGLERVSQDWTIKIEAGPGPAGSAVELAVRLTVFFITLIAPLVAGAVVYTKRRSSTQTETVPHGLEMTSHG-

**>Dicp5 (AMCP00006282, modified)**

MAVDSERGLLLLLLLLSNAAPLTAVVLYSTLGASVTVPCDGLTEYDNSYIVWVFAHSSDTSVELSRGGMITDTDPDRAGRLRLGSNRYLHIDRLRTRDTGEY
NCHQHVNGHFYESGSTVILFLLSISASPSEGLRAGDRITLSCVLDCGGMVGNCSETQGLTLSWRDESGFPLKDERDRYSITELRGVHSHLSVTLRQSDHNKS
RTCVLTERGEMKTSESYTTTLSDELFSTVGRLLLLPCVDPVSLGPGERLRWTYRQPVSSAPVTLYELLSQSITPLRGKDTHSGRAVMSVNSSLLIHNVQTRD
AGLYRCERHKDLKYQKTHRRFALNTLSVNSNPIADVHKGSAVSLTCSLTCGFDCVENTELTWKNSTGNSLQGGASEHNKSTISSQLVLQPQSSERIWCSVER
EGLERVSQDWTTEIKAGPGLAGSAVELAVRLTMFFIALIAPLVTGAVVYTKRRSSTQTEAVPQWMEMTSHG

**>Dicp6 (AMCP00006281, modified)**

MAVDSERGLLLLLLLLSTAAPLTDVTVFSTVGGSAALRCETVIYTDCSSTRWLFNRRSQAAVELVNLGKVKNNNLERAGRLSVGSDCSLHIDRLHTQDTGVY
TCRQFINEQKQGGDNTVYLALLIIEVPPETELKAGSTVTLRCLLHTGHGPVVCSHPPYTSADVRVSWVSETGAELQGDTYQISTDHPCLSTLTVRLQTSDHN
TQWRCDLTEGGAVRVSQRHTIKLTAGSAVELAVRLTVFFIALIAPLVAGVVVYTKTRSSRQTVPPLATALSLAMLLPVGVVIAVCVVIQRSRRQKVIEDVSD
PSAAADTVTFAVIDTSRARERDPATTGETEASNTEYATVRLH-

**>Dicp7 (partial)**

MAVDSERGLLLLLLLLNTAAPITGVAVYSSVGGSATLPCEGEINTDCSSSAWFFKETHMLDTNGKPGKLSVGSNCSLQIYNLSTEDAGNYSCHQQINGQHNN
KVNLALLSIEVSPETELKAGSSVTLLCLLNTDYGPAHCEQYPYVSADVSVNWVSEXXXXXXXXXXXXXXXXXXXX

**>Dicp9 (partial)**

MAVDSERGLLLLLLLSTAAPLTGVAVFSTVGGIADLHCKTVIYTNCSSTTWNFNSGSQTTVELVGLGKVKNNNPERAGRLSLGSNCSLHIDRLHTQDTGLYY
CQQFINGQKQGVDNTVYLALLI

**>Dicp10 (partial)**

MAVDRERGLLLLVMLSTAASLTGVAVFSTVGGIADLHCKTVIYTNCSSTVWNFNSGSQTTVELVGLGKVKNNNPERAGRLSLGSNCSLHIDRLHTQDTGLYY
CQQFINGKKQGVDYTVYLALLIIDVPPETELKAGSTVTLRCLLHTGHGPGDCSHPPYTSADVRLSWVSETGAELQGDTYFTERPCLSTLTVRLQTSDHNTQW
RCDLTEGGAVRVSQRHTIKLT

**>Dicp11 (AMCP00006287, modified)**

MAVDSERGLLLLLLLLNTAAPLTGVAVYSTVGGSATLPCEGVINTDCSSTTWLFIQKSQLAVELVELGKIRNYSPVRARRQSVGSDCSLYIDRLHTKDTGLY
YCQQFINGQKQGGDYSVYLVLLIIDVPPETELKAGSTVTLRCFLHTGHSPGVCSHPYTSADVRLSWVSETGAELQGDTYQISTDQPCLSTLTVRLQTSDHNT
QWRCDLTEGGAVRVSQRHTIKLTGIPEDNTTAATATAARTTTPKPNTTTTTKTGSHIETIVSLSIVLPVALLISAVAVYVGIRRSRRPRGNQDVADPSAQAA
DTVTYAVIETSKARERDPATAGGTEAPNTEYATVRLH

**>Dicp12 (AMCP00006283, modified)**

MRSPLFPLLISLSCLSEWAQRLGVEASVTVPCDGLTEYHNSYISWVFNHRSETTVELSRGGMITDTDPDRAGRLRLGSNSSLHIDRLRTRDTGQYNCHQYVN
GKYYTSGLTVTLFLLSITASPSENLKSGDRITLRCGLDCGGGAGSCSEAPQGLTLTWRDESGAPLKDRRDRYEIKTLANRSHLSVSLQRSDHYRLWVCALAE
GGRVETCGNYTTKLSDFIREVYIRLGDFLQLPCLESVHLGPGEILQWLVNKADTINYETLYTLSEDGQITTAMEVNPERLEMTVSSSLLIRSVQAQDSGVYL
CCFNEETHEVYYLNTIWVSSDHTGEVKRGSNITLTCTLTCGRIYGNISNTLVWRDSTGHSLQGGTTEQMNNKFISRLLVPELQSSERIWCSVVREGLERVEQ
DITIVVTGNSTLYDPQGSALSLVELAVRLTVFFIALFVPLVAVAVVYTKRRSSRQTGE-

**>Dicp14 (AMCP00006286, modified)**

MAVDSERGLLLLLLSTAAPLTGVAVFSTVGGSAALRCGTVIYTNCSSTTWNFNSGSQTTVELVGLGKVNNDSPERAGRLSVGSDCSLHIDRLHTQDTGHYYC
QQFINGQKQGGDYRVYLALLIIDVPPETELKAGSTVTLRCLLHTGHGPGVCSHPPYTSADVRVSWVSETGAELQGDTYQISTDHPCLSTLTVRLQTSDHNTQ
WRCDLTEGGAVRVSQRHTIKLTGSPIETIVSLSIILPVALLIAAVAVYVGIRRSSRPRGERPD-

**>Dicp16 (AMCP00006291, modified)**

VAVYSTVGGSATLPCEGATSGICSAVDWLFIKAPGELVHLVNHGRVTATDPERAGRLSVASDCSLHIDRLHTQDTGRYTCLPYFNGRDYLVNLALLSVPPET
ELKAGSIVTLRCFLDTEHGPGVCSHPYTSADVRLSWVSETGAELQGDTYQISTDQPCLSTLTVRLQTSDHNTQWRCDLTEGGAVRVSQRHTIKLT

**>Dicp19(AMCP00006292,modified)**
LSLYSSQGGEVTVSCEGLRKYENDSIAWSYSHGIGPPIELSRGMNITATDPGRAGRLRVLSDNSLHIDGLGPRDIGRYTCQQLINGRLNRSGSPVRLGLLYI
TTSPSENLKVNDTLTLTCTVSVEGSFPPKIPTLDWKIKRSTGRIIMTGVPGRINLSVKLHRSDHGRNWVCDLSEDGKIFSAIYTTKLTDEVYSTVGGLLLLP
CVDPVSLGRGETLRWTYRQPASSAPVTLYGLTSQSETPLAGKGVENQRAVMLANSSLLIHSVQTGDAGLYRCEENKTQEMLAVNILSVSSEQTGEVQKGNGS
AVTLACSLTCGSGCVEDTELIWKNSTGHSLQGSASECNKSTICSQLVLQPQSSERIRCSVEREGLERVSQDWTIKIEAGPGSAVELAVRLTVFFIALIAPLV
AGAVVYTKRRSSRQT

**>Dicp20(AMCP00006288)**
MAVDSERGLLLLLLLLSTAAPLTGVAVYSTVGGSATLPCEGATSGICSSIYWVFHKDDAELVGGGRVTASDPERAGRLSVKSDCSLHIDRLHTQDTGHYYCR
PHFNGDDYLVNLALLSVDVPPETELKAGSTVTLRCLLHTGHGPGDCSHPYTSADVRLSWVSETGAELQGDTYQISTDHPCLSTLTVRLQTSDHNTQWRCDLT
EGGAVRVSQRHTIKLT

**Fig. S1. Genome-predicted bowfin DICP proteins**.

DICP D1 domains are shaded dark orange and D2 domains are shaded light orange. Signal peptides and transmembrane (TM) domains are boxed. ITIM and ITIM-like sequences are in red text and a charged residue within a TM domain are shaded green. A frame shift (fs) and internal stop codon are indicated in Dicp21p. Pseudogenes and genes predicted with a single exon are not included. A run of 20 Xs (shaded gray) indicates a region where a DICP exon abuts an unresolved sequence in the reference genome.

**>"DICP3.3.1.1"**
MAVDSERGLLLLLLLSTAAPLTGVAVYSTVGGSATLPCEGATSGICSSIYWVFHKDDAELVGGGRVTASDPERAGRLSVKSDCSLHIDRLHTQDTGHYYCR
PHFNGDDYLVNLALLSVDVPPETELKAGSTVTLRCLLHTGHGPGDCSHPYTSADVRLSWVSETGAELQGDTYQISTDHPCLSTLTVRLQTSDHNTQWRCDLT
EGGAVRVSQRHTIKLTDVLSSGPSQLWIIGGTAVLLCVLGLVLGVWIWMSRRRRTRSGSEGHLMQCTVDSSNRREEEENHVKKDNKVGSENDCKFTDENYVS
IAFPPSKSGAKRKQHFRGKSKE**VVYTEV**KTRVRVSEGGN-

**>LOC100536667.1.1**
SVFNGKTTRSRGLSAASWQRQTRSPASAQQSRLASKCIELQHLFPCEGALTRQIQLRTKAEGVAVFSTVGGSAALRCGTVIYTNCSSTTWNFNSGSQTTVEL
VGLGKVNNDSPERAGRLSVGSDCSLHIDRLHTQDTGHYYCQQFINGQKQGGDYRVYLVLLIIDVPPETELKAGSTVTLRCLLHTGHSPGVCSHPPYTSADVR
LSWVSETGAELQGDTYFTERPCLSTLTVRLQTSDHNTQWRCDLTEGGAVRASQRHTIKLTGIPEDNTTPATTTATRTTTPKPNTTVTTKTGIPEDNTTAATA
TAARTTTPKPNTTTTTKTGSHIETIVSLSIVLPVALLISAVAVYVGIRRSRRPSTGNQDVADPSAAADT**VTYAVI**DTSRARERDPATAGGTEAPNTDAPQPS
RILTWTHLDSLPIFLTYAPDSLPGYFSWGFLSFSSSHTQITALDGVQWCFLGIKWMMWCWRGKWFKTAVQLRWQNHTGSRILINNNKKKKKKLGTRNGEHR
N-

**>LOC100333982.1.2**
MRSALFPLLISLSCLSEWAQRLGVEAVFLYSTLGASVTVPCDGLREYDNSYISWVFKHRSETTVELSRGGMITDTDPDRAGRLRLGSNSSLHIDRLRTRDTG
QYICHQYVNGMHYTTGLTVTLFLLSISASSRSSACLRGGDRLTLRCGLDCGGGAGSCSEAPQGLTLTWRDESGAPLKDRRDRYEIKTLANRSHLSVSLQRSD
HYRLWVCALAEGGRVETCGNYTTKLSDFIREVYIRLGDFLQLPCLESVHLGPGEILQWLVNKADTINYETLYTLSEDGQITTAMEVNPERLEMTVSSSLLIR
SVQAQDSGVYLCCFNEETHEVYYLNTIWVSSDHTGEVKRGSNITLTCTLTCGRIYGNISNTLVWRDSTGHSLQGGTTEQMNNKFISRLLVPELQSSERIWCS
VVREGLERVEQDITIVVTDPQGSALSLVELAVRLTVFFIALFVPLVAVAVVYTKRRSSRQTEEEETNIELVSRD-

**>LOC100333982.2.2**
MRSAIFPLLISLSCLSEWAQRLGVEAVFLYSTLGASVTVPCDGLREYDNSYISWVFKHRSETTVELSRGGMITDTDPDRAGRLRLGSNSSLHIDRLRTRDTG
QYICHQYVNGKHYTTGLTVTLFLLSISASSRSPACLRAGDRLTLKCGLDCGGGAGSCSETPQGPTLSWRDESGFPLKDERDRYSITELRGVHSQLSVTLRQS
DHNKSWTCVLTERGEMKTSESYTTTLSDELFSTVGRLLLLSCVAPVSLGPGETLRWTYRQSRTSQAVVLYSLRSLTETPLKGTVPHSGRAVMSANSSLLIHN
VQTGDAGLYCCQRDKDTGPKKTHRTFALNTLSVSSNLSEEAQKGSAVTLTCSLTCGFDCEENTELIWRDSIGNSLQGGTSERNKSTISSQLVLQPQSSERIR
CSVEREGLERVSQDWTIKIEAGPGGKFPMVEVVSAAAAFCVLLLLVFIIFLVMKMKKTKKEGKKDRVPSDQAEERPESVYHSPEEIMSCAAPPHTDD**VNYTS**
**V**MFKSKKRGPEREQTFLPNSDD**VIYSAV**TTTQ-

**>PTC1.1.2**
MITDTDPDRAGRLRLGSNSSLHIDRLRTRDTGQYTCHQYVNGKHYTTGLTVTLFLLSISASSRSPACLRAGDRLTLKCGLDCGGGAGSCSETPQGPTLSWRD
ESGFPLKDERDRYSITELRGVHSQLSVTLRQSDHNKSWTCVLTERGEMKTSESYTTTLSDELFSTVGRLLLLSCVAPVSLGPGETLRWTYRQSRTSQAVVLY
SLRSLTETPLKGTVPHSGRAVMSANSSLLIHNVQTGDAGLYCCQRDKDTGPKKTHRTFALNTLSVSSNLSEEAQKGSAVTLTCSLTCGFDCEENTELIWRDS
IGNSLQGGTSERNKSTISSQLVLQPQSSERIRCSVEREGLERVSQDWTIKIEAGPGPAGSAVELAVRLTVFFITLIVPLVAGAVVYTKRRSSTQTETVPHGL
EMTSHG

**>1382_c24_g1_i1**
MRSAIFPLLISLSCLSEWAQRLGVEAVFLYSTLGASVTVPCDGLREYHNSYISWVFKHRSETTVELSRGGMITDTDPDRAGRLRLGSNSSLHIDRLRTRDTG
QYTCHQYVNGKYYTSGLTVTLFLLSISASSRSPACLRAGDRLTLKCVLDCGGGAGSCSETPQGPTLSWRDESGFPLKDERDRYSITELRGVHSQLSVTLRQS
DHNKSWTCVLTERGEMKTSESYTTTLSDELFSTVGRLLLLSCVAPVSLGPGETLRWTYRQSRTSQAVVLYSLRSLTETPLKGTVPHSGRAVMSANSSLLIHN
VQTGDAGLYCCQRDKDSGPKKTHRTFALNTLSVSSNLSEEAQKGSAVTLTCSLTCGFDCEENTELIWRDSIGNSLQGGTSERNKSTISSQLVLQPQSSERIR
CSVEREGLERVSQDWTIKIEAGPGGKFPMVEVVSAAAAFCVLLLLVFIIFLVMKMKKTKKEGKKDRVPSDQAEERPESVYHSPEEIMSCAAPPHTDD**VNYTS**
**V**MFKSKKRGPEREQTFLPNSDD**VIYSAV**TTTQ-

**>1382_c24_g1_i5**
MRSAIFPLLISLSCLSEWAQRLGVEAVFLYSTLGASVTVPCDGLREYHNSYISWVFKHRSETTVELSRGGMITDTDPDRAGRLRLGSNSSLHIDRLRTRDTG
QYTCHQYVNGKYYTSGLTVTLFLLSISASSRSPACLRAGDRLTLKCVLDCGGGAGSCSETPQGPTLSWRDESGFPLKDERDRYSITELRGVHSQLSVTLRQS
DHNKSWTCVLTERGEMKTSESYTTTLSDELFSTVGRLLLLSCVAPVSLGPGETLRWTYRQSRTSQAVVLYSLRSLTETPLKGTVPHSGRAVMSANSSLLIHN
VQTGDAGLYCCQRDKDSGPKKTHRTFALNTLSVSSNLSEEAQKGSAVTLTCSLTCGFDCEENTELIWRDSIGNSLQGGTSERNKSTISSQLVLQPQSSERIR
CSVEREGLERVSQDWTIKIEAGPGEELDL

**>1382_c35_g3_i1**
MAVDSERGLLLLLLLSTAASLTGVAVFSTVGGIADLHCKTVIYTNCSSTTWNFNSGSQTTVELVGLGKVKNNNPERAGRLSVGSNCSLHIDRLHTQDTGLYY
CQQFINGKKQGVDNTVYLVLLIIDVPPETELKAGSTVTLRCLLHTGHSPGVCSHPPYTSADVRLSWVSETGAELQGDTYFTERPCLSTLTVRLQTSDHNTQW
RCDLTEGGAVRVSQRHTIKLTDVLSSGPSQLWIIGGTAVLLCVLGLVLGVWIWMSRRRRTRSGSEGHLMQCTVDSSNRREEEENHVKKDNKVGSENDCKFTD
**ENYVSI**AFPPSKSGAKRKQHFRGKSKEVVYTE

**>1382_c35_g3_i5**
MAVDSERGLLLLLLLSTAASLTGVAVFSTVGGIADLHCKTVIYTNCSSTTWNFNSGSQTTVELVGLGKVKNNNPERAGRLSVGSNCSLHIDRLHTQDTGLYY
CQQFINGKKQGVDNTVYLVLLIIDVPPETELKAGSTVTLRCLLHTGHSPGVCSHPPYTSADVRLSWVSETGAELQGDTYFTERPCLSTLTVRLQTSDHNTQW
RCDLTEGGAVRVSQRHTIKLTGIPEDNTTTATTATRTTTPKPNTTVTTKTGSAVELVVRLTVFFIALIAPLVAVAVVYSKRRSSRQTEEVPSGIELDSTG-

**Fig. S2. Transcriptome-predicted bowfin DICP proteins**.

DICP D1 domains are shaded dark orange and D2 domains are shaded light orange. Signal peptides and transmembrane (TM) domains are boxed. ITIM and ITIM-like sequences are in red text and a charged residue within a TM domain are shaded green.

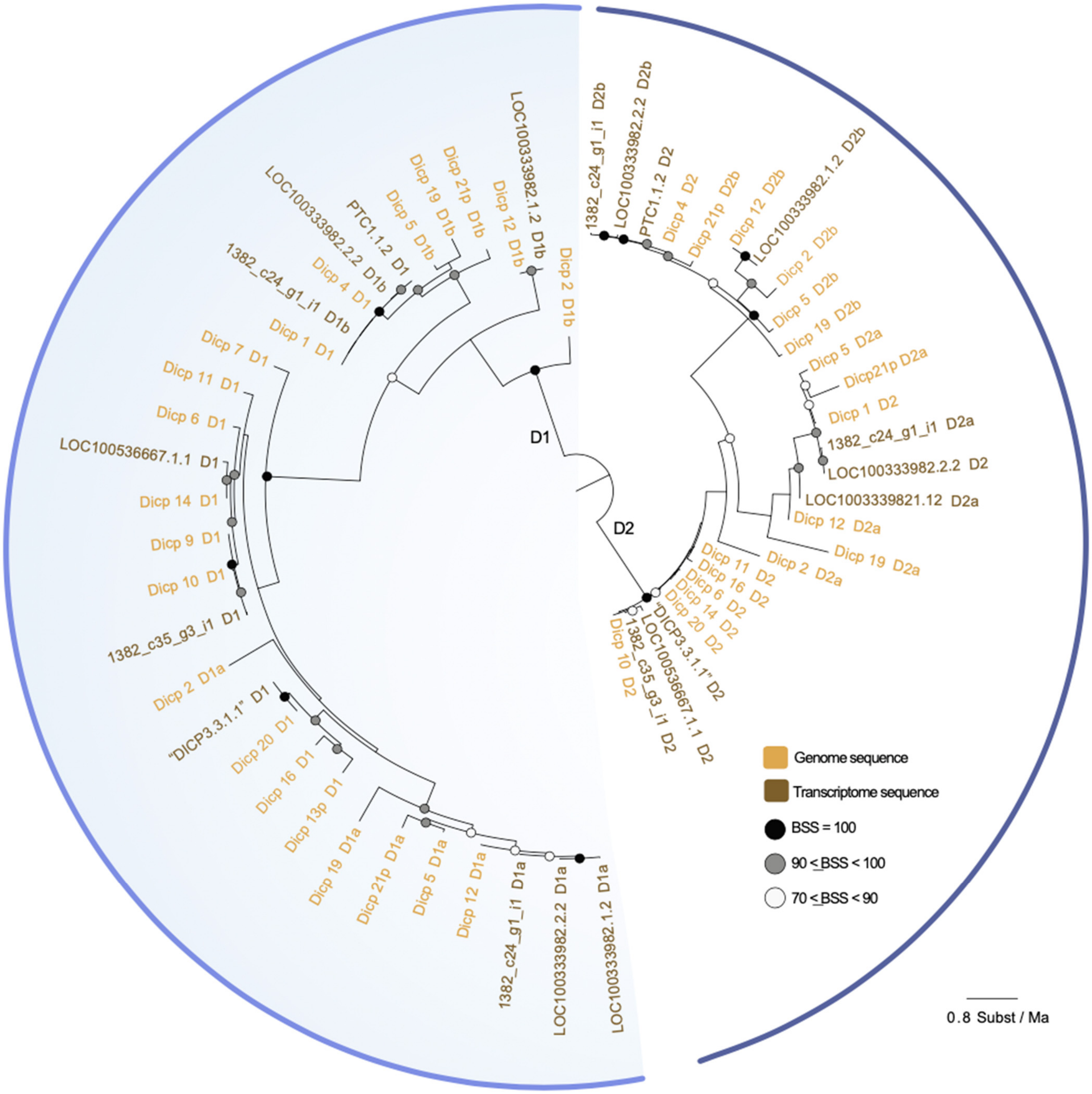

**Fig. S3. Bowfin DICP transcripts do not match (exactly) the reference genome**.
The phylogenetic relationships of bowfin DICP D1 and D2 domains identified from the bowfin genome (light brown) and transcripts (dark brown) inferred using maximum likelihood in IQ-TREE. Circles at nodes indicate bootstrap support values (BSS) with filled black circles black indicating BSS=100, gray circles indicating BSS values equal to or greater than 90 but less than 100, and white circles indicating BSS values greater than 70 but less than 90. D1 and D2 lineages are indicated by the color coded lines outside of the circle (light blue = D1; dark blue = D2).

```
Dicp20            1 MAVDSERGLLLLLLLLSTAAPLTGVAVYSTVGGSATLPCEGATSGICSSIYWVFHKDDAELVGGGRVTASDPERAGRLSV
"DICP3.3.1.1"     1 MAVDSERGLLLLLLLLSTAAPLTGVAVYSTVGGSATLPCEGATSGICSSIYWVFHKDDAELVGGGRVTASDPERAGRLSV

Dicp20           81 KSDCSLHIDRLHTQDTGHYYCRPHFNGDDYLVNLALLSVDVPPETELKAGSTVTLRCLLHTGHGPGDCSHPYTSADVRLS
"DICP3.3.1.1"    81 KSDCSLHIDRLHTQDTGHYYCRPHFNGDDYLVNLALLSVDVPPETELKAGSTVTLRCLLHTGHGPGDCSHPYTSADVRLS

Dicp20          161 WVSETGAELQGDTYQISTDHPCLSTLTVRLQTSDHNTQWRCDLTEGGAVRVSQRHTIKLT--------------------
"DICP3.3.1.1"   161 WVSETGAELQGDTYQISTDHPCLSTLTVRLQTSDHNTQWRCDLTEGGAVRVSQRHTIKLTDVLSSGPSQLWIIGGTAVLL

Dicp20              --------------------------------------------------------------------------------
"DICP3.3.1.1"   241 CVLGLVLGVWIWMSRRRRTRSGSEGHLMQCTVDSSNRREEEENHVKKDNKVGSENDCKFTDENYVSIAFPPSKSGAKRKQ

Dicp20              -------------------------
"DICP3.3.1.1"   321 HFRGKSKEVVYTEVKTRVRVSEGGN
```

**Fig. S4. Dicp20 and "DICP3.3.1.1".**

Alignment of bowfin proteins encoded by the *dicp20* gene and *"DICP3.3.1.1"* transcript.

```
Dicp1(partial)      1 MRSAIFPLLISLSCLSEWAQRLGVEXXXXXXXXXX------------------------------------------XXX
1382_c24_g1_i1      1 MRSAIFPLLISLSCLSEWAQRLGVEAVFLYSTLGASVTVPCDGLREYHNSYISWVFKHRSETTVELSRGGMITDTDPDRA
LOC100333982.2.2    1 MRSAIFPLLISLSCLSEWAQRLGVEAVFLYSTLGASVTVPCDGLREYDNSYISWVFKHRSETTVELSRGGMITDTDPDRA

Dicp1(partial)     39 XXXXXXXNSSLHIDRLRTRDTGQYTCHQYVNGKHYTSGLPVTLFLLSISVSSSSSACLRAGDRLTLKCGLDCGGGAGSCS
1382_c24_g1_i1     81 GRLRLGSNSSLHIDRLRTRDTGQYTCHQYVNGKYYTSGLTVTLFLLSISASSRSPACLRAGDRLTLKCVLDCGGGAGSCS
LOC100333982.2.2   81 GRLRLGSNSSLHIDRLRTRDTGQYICHQYVNGKHYTTGLTVTLFLLSISASSRSPACLRAGDRLTLKCGLDCGGGAGSCS

Dicp1(partial)    119 ETPQGLTLSWRDESGVPPKDERDRYSITELRGVHSQLSVTLRQSDHNKSWTCVLTERGEMKTSESYTTTLSDELFSTVGR
1382_c24_g1_i1    161 ETPQGPTLSWRDESGFPLKDERDRYSITELRGVHSQLSVTLRQSDHNKSWTCVLTERGEMKTSESYTTTLSDELFSTVGR
LOC100333982.2.2  161 ETPQGPTLSWRDESGFPLKDERDRYSITELRGVHSQLSVTLRQSDHNKSWTCVLTERGEMKTSESYTTTLSDELFSTVGR

Dicp1(partial)    199 LLLLSCVAPVSLGPGETLRWTYRQSRTSQAVVLYSLRSLTETPLKGTVPHSGRAVMSANSSLLIHNVQTGDAGLYCCQRD
1382_c24_g1_i1    241 LLLLSCVAPVSLGPGETLRWTYRQSRTSQAVVLYSLRSLTETPLKGTVPHSGRAVMSANSSLLIHNVQTGDAGLYCCQRD
LOC100333982.2.2  241 LLLLSCVAPVSLGPGETLRWTYRQSRTSQAVVLYSLRSLTETPLKGTVPHSGRAVMSANSSLLIHNVQTGDAGLYCCQRD

Dicp1(partial)    279 KDSGPKKTHRTFALNTLSXXXXXXXXXX-------------------------------------XXXXXXXXXXQLVLQ
1382_c24_g1_i1    321 KDSGPKKTHRTFALNTLSVSSNLSEEAQKGSAVTLTCSLTCGFDCEENTELIWRDSIGNSLQGGTSERNKSTISSQLVLQ
LOC100333982.2.2  321 KDTGPKKTHRTFALNTLSVSSNLSEEAQKGSAVTLTCSLTCGFDCEENTELIWRDSIGNSLQGGTSERNKSTISSQLVLQ

Dicp1(partial)    322 PQSSERIRCSVERQGLERVSQDLTIKIEAGPGGKFPMVEVVSAAAAFCVLLLLVFIIFLVMKMKKTKKEGKKDRVPSDQA
1382_c24_g1_i1    401 PQSSERIRCSVEREGLERVSQDWTIKIEAGPGGKFPMVEVVSAAAAFCVLLLLVFIIFLVMKMKKTKKEGKKDRVPSDQA
LOC100333982.2.2  401 PQSSERIRCSVEREGLERVSQDWTIKIEAGPGGKFPMVEVVSAAAAFCVLLLLVFIIFLVMKMKKTKKEGKKDRVPSDQA

Dicp1(partial)    402 EERPESVYHSPEEIMSCAAPPHTDDVNYTSVMFKSKKRGPEREQTFLPNSDDVIYSAVTTTQ
1382_c24_g1_i1    481 EERPESVYHSPEEIMSCAAPPHTDDVNYTSVMFKSKKRGPEREQTFLPNSDDVIYSAVTTTQ
LOC100333982.2.2  481 EERPESVYHSPEEIMSCAAPPHTDDVNYTSVMFKSKKRGPEREQTFLPNSDDVIYSAVTTTQ
```

**Fig. S5. Dicp1, 1382_c24_g1_i1 and LOC100333982.2.2.**

Alignment of bowfin proteins encoded by the *dicp1* gene and *1382_c24_g1_i1 and LOC100333982.2.2* transcripts.

```
Dicp12            1 MRSPLFPLLISLSCLSEWAQRLGVEA---------SVTVPCDGLTEYHNSYISWVFNHRSETTVELSRGGMITDTDPDRA
LOC100333982.1.2  1 MRSALFPLLISLSCLSEWAQRLGVEAVFLYSTLGASVTVPCDGLREYDNSYISWVFKHRSETTVELSRGGMITDTDPDRA

Dicp12           72 GRLRLGSNSSLHIDRLRTRDTGQYNCHQYVNGKYYTSGLTVTLFLLSITASP--SENLKSGDRITLRCGLDCGGGAGSCS
LOC100333982.1.2 81 GRLRLGSNSSLHIDRLRTRDTGQYICHQYVNGMHYTTGLTVTLFLLSISASSRSSACLRGGDRLTLRCGLDCGGGAGSCS

Dicp12          150 EAPQGLTLTWRDESGAPLKDRRDRYEIKTLANRSHLSVSLQRSDHYRLWVCALAEGGRVETCGNYTTKLSDFIREVYIRL
LOC100333982.1.2 161 EAPQGLTLTWRDESGAPLKDRRDRYEIKTLANRSHLSVSLQRSDHYRLWVCALAEGGRVETCGNYTTKLSDFIREVYIRL

Dicp12          230 GDFLQLPCLESVHLGPGEILQWLVNKADTINYETLYTLSEDGQITTAMEVNPERLEMTVSSSLLIRSVQAQDSGVYLCCF
LOC100333982.1.2 241 GDFLQLPCLESVHLGPGEILQWLVNKADTINYETLYTLSEDGQITTAMEVNPERLEMTVSSSLLIRSVQAQDSGVYLCCF

Dicp12          310 NEETHEVYYLNTIWVSSDHTGEVKRGSNITLTCTLTCGRIYGNISNTLVWRDSTGHSLQGGTTEQMNNKFISRLLVPELQ
LOC100333982.1.2 321 NEETHEVYYLNTIWVSSDHTGEVKRGSNITLTCTLTCGRIYGNISNTLVWRDSTGHSLQGGTTEQMNNKFISRLLVPELQ

Dicp12          390 SSERIWCSVVREGLERVEQDITIVVTGNSTLYDPQGSALSLVELAVRLTVFFIALFVPLVAVAVVYTKRRSSRQTGE---
LOC100333982.1.2 401 SSERIWCSVVREGLERVEQDITIVVT------DPQGSALSLVELAVRLTVFFIALFVPLVAVAVVYTKRRSSRQTEEEET

Dicp12              --------
LOC100333982.1.2 475 NIELVSRD
```

**Fig. S6. Dicp12 and LOC100333982.1.2.**

Alignment of bowfin proteins encoded by the *dicp12* gene and *LOC100333982.1.2* transcript.

```
Dicp14             1 ----------------------------------------MAVDSERGLLLLLLSTAAPLTGVAVFSTVGGSAALRCGTV
LOC100536667.1.1   1 SVFNGKTTRSRGLSAASWQRQTRSPASAQQSRLASKCIELQHLFPCEGALTRQIQLRTKAEGVAVFSTVGGSAALRCGTV

Dicp14            41 IYTNCSSTTWNFNSGSQTTVELVGLGKVNNDSPERAGRLSVGSDCSLHIDRLHTQDTGHYYCQQFINGQKQGGDYRVYLA
LOC100536667.1.1  81 IYTNCSSTTWNFNSGSQTTVELVGLGKVNNDSPERAGRLSVGSDCSLHIDRLHTQDTGHYYCQQFINGQKQGGDYRVYLV

Dicp14           121 LLIIDVPPETELKAGSTVTLRCLLHTGHGPGVCSHPPYTSADVRVSWVSETGAELQGDTYQISTDHPCLSTLTVRLQTSD
LOC100536667.1.1 161 LLIIDVPPETELKAGSTVTLRCLLHTGHSPGVCSHPPYTSADVRLSWVSETGAELQGDTYF--TERPCLSTLTVRLQTSD

Dicp14           201 HNTQWRCDLTEGGAVRVSQRHTIKLTGS----------------------------------------------------
LOC100536667.1.1 239 HNTQWRCDLTEGGAVRASQRHTIKLTGIPEDNTTPATTTATRTTTPKPNTTVTTKTGIPEDNTTAATATAARTTTPKPNT

Dicp14           229 --------PIETIVSLSIILPVALLIAAVAVYVGIRRSSRPRGERPD*--------------------------------
LOC100536667.1.1 319 TTTTKTGSHIETIVSLSIVLPVALLISAVAVYVGIRRSRRPSTGNQDVADPSAAADTVTYAVIDTSRARERDPATAGGTE

Dicp14               --------------------------------------------------------------------------------
LOC100536667.1.1 399 APNTDAPQPSRILTWTHLDSLPIFLTYAPDSLPGYFSWGFLSFSSSHTQITALDGVQWCFLGIKWMMWCWRGKWFKTAVQ

Dicp14               ---------------------------------
LOC100536667.1.1 479 LRWQNHTGSRILINNNKKKKKKKLGTRNGEHRN
```

**Fig. S7. Dicp14 and LOC100536667.1.1.**

Alignment of bowfin proteins encoded by the *dicp14* gene and *LOC100536667.1.1* transcript.

```
Dicp9(partial)     1 MAVDSERGLLLLLLLSTAAPLTGVAVFSTVGGIADLHCKTVIYTNCSSTTWNFNSGSQTTVELVGLGKVKNNNPERAGRL
Dicp10(partial)    1 MAVDRERGLLLLVMLSTAASLTGVAVFSTVGGIADLHCKTVIYTNCSSTVWNFNSGSQTTVELVGLGKVKNNNPERAGRL
1382_c35_g3_i1     1 MAVDSERGLLLLLLLSTAASLTGVAVFSTVGGIADLHCKTVIYTNCSSTTWNFNSGSQTTVELVGLGKVKNNNPERAGRL
1382_c35_g3_i5     1 MAVDSERGLLLLLLLSTAASLTGVAVFSTVGGIADLHCKTVIYTNCSSTTWNFNSGSQTTVELVGLGKVKNNNPERAGRL

Dicp9(partial)    81 SLGSNCSLHIDRLHTQDTGLYYCQQFINGQKQGVDNTVYLALLI------------------------------------
Dicp10(partial)   81 SLGSNCSLHIDRLHTQDTGLYYCQQFINGKKQGVDYTVYLALLIIDVPPETELKAGSTVTLRCLLHTGHGPGDCSHPPYT
1382_c35_g3_i1    81 SVGSNCSLHIDRLHTQDTGLYYCQQFINGKKQGVDNTVYLVLLIIDVPPETELKAGSTVTLRCLLHTGHSPGVCSHPPYT
1382_c35_g3_i5    81 SVGSNCSLHIDRLHTQDTGLYYCQQFINGKKQGVDNTVYLVLLIIDVPPETELKAGSTVTLRCLLHTGHSPGVCSHPPYT

Dicp9(partial)       --------------------------------------------------------------------------------
Dicp10(partial   161 SADVRLSWVSETGAELQGDTYFTERPCLSTLTVRLQTSDHNTQWRCDLTEGGAVRVSQRHTIKLT---------------
1382_c35_g3_i1   161 SADVRLSWVSETGAELQGDTYFTERPCLSTLTVRLQTSDHNTQWRCDLTEGGAVRVSQRHTIKLTDVLSSGPSQLWI---
1382_c35_g3_i5   161 SADVRLSWVSETGAELQGDTYFTERPCLSTLTVRLQTSDHNTQWRCDLTEGGAVRVSQRHTIKLTGIPEDNTTTATTATR

Dicp9(partial)       --------------------------------------------------------------------------------
Dicp10(partial)      --------------------------------------------------------------------------------
1382_c35_g3_i1   238 ------------IGGTAV---------LLCVLGLVLGVWIWMSRRRRTRSGSEGHLMQCTVDSSNRREEEENHVKKDNKV
1382_c35_g3_i5   241 TTTPKPNTTVTTKTGSAVELVVRLTVFFIALIAPLVAVAVVYSKRRSSRQTEE-VPSGIELDSTG---------------

Dicp9(partial)       ------------------------------------------
Dicp10(partial)      ------------------------------------------
1382_c35_g3_i1   297 GSENDCKFTDENYVSIAFPPSKSGAKRKQHFRGKSKEVVYTE
1382_c35_g3_i5       ------------------------------------------
```

**Fig. S8. Dicp9, Dicp10, 1382_c35_g3_i1 and 1382_c35_g3_i5.**

Alignment of bowfin proteins encoded by the *dicp9* and *dicp10* genes and by the *1382_c35_g3_i1* and *1382_c35_g3_i5* transcripts.

```
1382_c24_g1_i1     1 MRSAIFPLLISLSCLSEWAQRLGVEAVFLYSTLGASVTVPCDGLREYHNSYISWVFKHRSETTVELSRGGMITDTDPDRA
1382_c24_g1_i5     1 MRSAIFPLLISLSCLSEWAQRLGVEAVFLYSTLGASVTVPCDGLREYHNSYISWVFKHRSETTVELSRGGMITDTDPDRA

1382_c24_g1_i1    81 GRLRLGSNSSLHIDRLRTRDTGQYTCHQYVNGKYYTSGLTVTLFLLSISASSRSPACLRAGDRLTLKCVLDCGGGAGSCS
1382_c24_g1_i5    81 GRLRLGSNSSLHIDRLRTRDTGQYTCHQYVNGKYYTSGLTVTLFLLSISASSRSPACLRAGDRLTLKCVLDCGGGAGSCS

1382_c24_g1_i1   161 ETPQGPTLSWRDESGFPLKDERDRYSITELRGVHSQLSVTLRQSDHNKSWTCVLTERGEMKTSESYTTTLSDELFSTVGR
1382_c24_g1_i5   161 ETPQGPTLSWRDESGFPLKDERDRYSITELRGVHSQLSVTLRQSDHNKSWTCVLTERGEMKTSESYTTTLSDELFSTVGR

1382_c24_g1_i1   241 LLLLSCVAPVSLGPGETLRWTYRQSRTSQAVVLYSLRSLTETPLKGTVPHSGRAVMSANSSLLIHNVQTGDAGLYCCQRD
1382_c24_g1_i5   241 LLLLSCVAPVSLGPGETLRWTYRQSRTSQAVVLYSLRSLTETPLKGTVPHSGRAVMSANSSLLIHNVQTGDAGLYCCQRD

1382_c24_g1_i1   321 KDSGPKKTHRTFALNTLSVSSNLSEEAQKGSAVTLTCSLTCGFDCEENTELIWRDSIGNSLQGGTSERNKSTISSQLVLQ
1382_c24_g1_i5   321 KDSGPKKTHRTFALNTLSVSSNLSEEAQKGSAVTLTCSLTCGFDCEENTELIWRDSIGNSLQGGTSERNKSTISSQLVLQ

1382_c24_g1_i1   401 PQSSERIRCSVEREGLERVSQDWTIKIEAGPGGKFPMVEVVSAAAAFCVLLLLVFIIFLVMKMKKTKKEGKKDRVPSDQA
1382_c24_g1_i5   401 PQSSERIRCSVEREGLERVSQDWTIKIEAGPGEELDL-------------------------------------------

1382_c24_g1_i1   481 EERPESVYHSPEEIMSCAAPPHTDDVNYTSVMFKSKKRGPEREQTFLPNSDDVIYSAVTTTQ
1382_c24_g1_i5       --------------------------------------------------------------
```

**Fig. S9. DICP transcripts 1382_c24_g1_i1 and 1382_c24_g1_i5.**

Alignment of bowfin proteins encoded by the *1382_c24_g1_i1* and *1382_c24_g1_i5* transcripts.

**>Nitr2**
GGSVRLSCYIRREASISTLTWIKQKPGDPPRPIGSWENNVASLQDEFKNSKRFAIERSGDDFDLKISSTDRSDVGRYFCAAAQSRSVRFGQGTVIQLPPVSV
PVQPRDNVTLQCVTDYYTCKGEHSVYWFRHRSGESLSGLITHKNQIDQCERSSELQFSTEYCVCSLTKTNLSSSDAGTYYCAVATSGEILFGNGTRLDIADR
SEAGGSAEGPCGCGGHSALSLSVLALAAVLCVFGT

**>Nitr16**
MIRLCVTLLLFCKVVNEGTVRSTVLRTAQLGDSVTLLCDLQGSYLIWFKQTTAQTPRSIATSYNYLPEITLFNEFKTDARFTVKRDEEGFNLTIARTEPSDE
ATYFCGILRTNHVHFGNGTYLTLKSGSKSVSRRVEQQPASVPVQSGDSVTLQCTIYTEPCAGEHSVHWFRHGSGEALPGLIYTHGNRSDPCESSSEPGLPAQ
GCVYELPKRNLCSSDAGTYYCAVATCGQILFGNGTRMDIA

**>Nitr20**
FPSHDGFVSDMTQTAATTLPFFFKTVQRGDSVTLGCLLATDRISYKAWFKQSTGFIPQVIAYSDMYLEKATFHNEFKDDPRFAVKADNHFQLTISGAEPSDS
AAYYCGSLYLKKWQFGNGTFVMVRVTSESESVIRRVEQRPASVLVQPGDSVTLQCTIHTETCAGEHSVHWFRHGSGEALPGLIYTHGNRSDLCESSSEPGLP
AQSCVYELPKRNLRSSDAGTYYCAVATCGQILFGNGTRLDIAGDESKGIFTLAVLALVTLNFISVIVISVLVCKLHQHTKCEYGA

**>Nitr21**
MIRLCVTLLLFCQVLAQTENVVQPRLYTAAQLGDSVTLECFLPSQKSTYTVWFKQTIGQRPQCMATAYNYLQETTFYEEFKHNPRFTVQRERDSFHLNISRT
ELSDTATYYCGVMFLNHMQFGNGTFLMVKISGSVSRRVEQQPASVPVQPGDSVTLQCTIHTETCAGEHSVHWFRHGSGEALPGLIYTHGNRSDPCESSSEPG
LPAQGCVYELPKRNLRSSDAGTYYCAVATCGQILFGNGTRLEFAG

**>Nitr24**
SCLVPDSVQMEDVIQPSLFAAAQPGDSVTLECYVPSDKVSYMSWFKQTIGQKPRPIAVSYAHVSDATFLDEFKGNPRFKVHTEDTQYHLTISRTEPSDTATY
YCVTMYTNVVNFGSGTFLLVKGKGSGSVSRRVEQQPASVPVQPGDSVTLQCTIHTETCAGEHSVHWFRHGSGEALPGLIYTHGNRSDPFESSSEPGLPAQGC
VYELPKRNLRSSDAGTYYCAVATCGQILFGNGTRLEFAG

**>Nitr27(AMCP00014119)**
LVLSQFYIQTPPSVTVDRGGSVRLSCYIRREASVSTLTWIKQEPGNPPRPIGSWENNADSLQDEFKNSSRFAIERSGDDFDLKISSTERSDVGRYFCAAAQS
RSVRFGQGTVIQMKDLDPESISRRAVQPPVSVPVQPGDNVTLQCVIDYYTCKAEHSVDWFRHRSGESLSGLKTHNNWIDQCDRSEAGGSAESPCGCGGHSAL
SLSVLALAAVLCVFGTLVTALLCRRHRAEAASGRPPHSDQHCSPAHGTDTESQLFV**LLYIRG**RHPGLSPLTKIHSTREICERDGQWSLLTLSHRVLSTSSYR
DGC

**>Nitr28**
TVELGGSVTLSCNVSKSNSADVLVWNKQALGKLSTSIASFQNNEASFHFEFYNNKHFAIDRRGDSFNLNISNIEASDAARYYCGAIRETSVRFGSGTSIRLK
SGSVSRRVEQQPASVPVQPGDSVTLQCTIHTETCAGEHSVHWFRHGSGEALPGLIYTHGNRSDPCESSSEPGLPAQGCVYELPKRNLRSSDAGTYYCAVATC
GQILFGNGARLHIADRSEAGGSAEGPCGCGGHSALSLSVLALAAVLCVFGTLVTALLCRRHRAGTGPQGAGR**LHYTAL**HMTVLRVCSITVSITNKWDVHELQ
YNSTL

**>Nitr34**
MNIILFLLSFCSKALLLCQLFAQSPSSVTVKLGGSVTLSCSVSKSYSSNVLIWNKQPSGGPSTSIVSFKNHESPLLGGDNKNKSFTIDRKGDSLNLKISNIE
ASDVARYYCGAIRDTSVRFGSGTSIRLKGSGSVSRRVEQQPASVPVQPGDSVTLQCTIHTETCAGEHSVHWFRHGSGEALPGLIYTHGNRSDPCESSSEPGL
PAQGCVYELPKRNLRSSDAGTYYCAVATCGPILFGNGTRLDIAGKKR

**Fig. S10. Genome-predicted bowfin NITR proteins.**
NITR V domains are shaded dark blue and I domains are shaded light blue. Signal peptides and transmembrane domains are boxed. ITIM and ITIM-like (itim) sequences are in red text. Pseudogenes and genes predicted from a single exon or with a single Ig domain are not included, with the exception of Nitr27 that encodes a truncated I domain.

**>"NITR1.1.2"**
MNIILFLLSFCSKAGLLLCQLFAQSPSSVTVELGGSVTLSCSVSKSYSSNVLFWNKQPSGGPSTSIVSFKNNAASFHGGSDNKRCAIDRRGHYFNLKISNIN
ASDVARYYCGAIRDTTVRFGSGTSIRLKGSGSVSRRVEQQPASVPVQPGDSVTLQCTIHTETCAGEHSVHWFRHGSGEALPGLIYTHGNRSDPCESSSEPGL
PAQGCVYELPKRNLRSSDAGTYYCAVATCGQILFGNGTRLDIAGDRSEAGGSAEGPCGCGGHSALSLSVLALAAVLCVFGTLVTALLCRRHRAGTGPQGAES
GHRADELRCPEVHCQQSSASWEKEARDGPTCAVC*****

**>"NITR1.2.2"**
MNIILFLLSFCSKAGLLLCQLFAQSPSSVTVELGGSVTLSCSVSKSYSSNVLFWNKQPSGGPSTSIVSFKNNAASFHGGSDNKRCAIDRRGHYFNLKISNIN
ASDVARYYCGAIRDTAVRFGSGTSIRLKGSGSVSRRVEQQPASVPVQPGDSVTLQCTIHTETCAGEHSVHWFRHGSGEALPGLIYTHGNRSDPCESSSEPGL
PAQGCVYELPKRNLRSSDAGTYYCAVATCGQILFGNGTRLDIAESSLQLSPAVLVLLASNCVCVLVIAVLVCTRRGQCGHGAEHVSQQTPHRHTGADWADKQ
NQGTGM**MNYAAL**SFTDSRTKPERKRREMDRQ**AVYSQV**QYHQTD*

**>"NITR3.1.2"**
KQPSGGPSATTVSFKNHETSFYLGFNNKKRFAIDRRGDSFNLKISNIEASDVARYYCGAIRDTTVRFGSGTSIRLKGSGSVSRRVEQQPASVPVQPGDSVTL
QCTIHTETCAGEHSVHWFRHGSGEALPGLIYTHGNRSDPCESSSEPGLPAQGCVYELPKRNLRSSDAGTYYCAVATCGQILFGNGTRLDIAAGSWLAGAGVV
VVWALAGLGLVSLALLAVLAFRVWSRHRAARRADLCQIAEDFSLPTSAETTSSQDAE**VTYTTV**ELGQRGRKHKQRRREQEQVDG**VVYTDI**RYHHRK*

**>"NITR3.2.2"**
MSSSLPDSLCFPGLLLCQLFAQSPSSVTVELGGSVTLSCSVSKSYSSNALVWIKQPSGGPSTSIVSFKNNAALFLGGFDNNKRCAIDRRGDSFNLNISNIEA
SDVARYYCGAIRDTTVRFGSGTSIRLKGSGSVSRRVEQQPASVPVQPGDSVTLQCTIHTETCAGEHSVHWFRHGSGEALPGLIYTHGNRSDPCESSSEPGLP
AQGCVYELPKRNLRSSDAGTYYCAVATCGQILFGNGTRLEFAGDRSEAGGSAEGPCGCGGHSALSLSVLALAAVLCVFGTLVTALLCRRHRAGTGPQGAEAA
SGRPPHSDQHCSPAHDTSTESQRGERDV**LTYAAL**DFDQNKRKAGKKRREGRSGSELV**SEYAAV**RTQRR*

**>"NITR4.1.2"**
*LFLFIGLDFLPLLTLPCAPHVGPPLWRPQSCISVVWVHAQPLIKSCRLWCSAVGVLCLPGVLLTDVVSQPQLSVTAQLRDSVTLHCSVFKTHPMTLAWLKQ
DVGQMPRYIATLSKYSQEMELQDEFKEGRFTVQNTSKSFDLQISHIRSSDVAVYYCAAFMYRRVACGNGTLLLLKVSGSVSRRVEQQPASVPVQPGDSVTLQ
CTIHTETCAGEHSVHWFRHGSGEALPGLIYTHGNRSDPCESSSEPGLPAQGCVYELPKRNLRSSDAGTYYCAVATCGQILFGNGTRLEFAAGSWLAGAGVVV
VWALAGLGLVSLALLAVLAFRVWSRHRAARRADLCQIAEDFSLPTSAETTSSQDAE**VTYTTV**ELGQRGRKHKQRRREQEQVDG**VVYTDI**RYHHRK*

**>"NITR4.2.2"**
MIRLCLPLCLLRTSVILADAITQPGFLLRAQLGDSVSLQCFIHQQSLKQLFWYKQVVGEAPKCMFSSYGQSGQLTRHGEFNHSRFSVQRTGSAFHLCISNTE
PTDIGTYYCATLDYSQVLFGGGSTLLLRDTDSRFIQPPLFQSLQSGLNSDLQCMMESRNLSADHRVHWFIKLSGQPSSKTPFTAGSMIQSSSEDDLPAQGCV
YNLSKRNLSCSDAESVYCAVATCGEILFGKRTEQEVTGRGVGVSHVPVVLLGALNAVLVVVIAVLLCRRQRSVQFQCCAARESGPVSQSPPQRRLTDSTSLQ
NQTIPT**VTYAAL**DFHHIKSKQERSRQDLQG**TVYRYI**LDQQWE*

**>LOC100695950.2.2**
MNIILFLLSFCSKAGLLLCQLFAQSPSSVTVELGGSVTLSCSVSKSYSSNVLVWNKQPSGGPSTSIVSFKNNAALFHGGFDNNKRCAIDRRGHYFNLKISNI
EASDVARYYCGAIRDTTVRFGSGTSIRLKGSGSVSRRVEQQPASVPVQPGDSVTLQCTIHTETCAGEHSVHWFRHGSGEALPGLIYTHGNRSDPCESSSEPG
LPAQGCVYELPKRNLRSSDAGTYYCAVATCGQILFGNGTRLDIAGDRSEAGGSVEGPSGCGGHSALSLSVLALAAVLCVFGTLVTALLCRRRRAGMGPQGAW
R**LLYTAL**RMTVLRVCSITVSITNKWDAHELQYDSSL*

**>4380_c1_g1_i1**
MEILQTKNRSFNLTILSTKPSDVAAYYCGAVYFNYMRFGKGTILILKDSESVIRRVEQQPASVPVQPGDSVTLQCTIHTETCAGEHSVHWFRHGSGEALPGL
IYTHGNRSDPCESSSEPGLPAQGCVYELPMRNLRSSDAGTYYCAVATCGQILFGNGTQLDIA

**>4857_c0_g1_i1**
GLLLCQLFAQSPSSVTVELGGSVTLSCSVSKSYSSNVLFWNKQPSGEPSTSIVSFTNNAASFHGGSDNKRCAIDRRGHYFNLKISNIEASDVARYYCGAIRD
TTVRFGSGTSIRLKGSGSVSRRVEQQPASVPVQPGDSVTLQCTIHTETCAGEHSVHWFRHGSGEALPGLIYTHGNRSDPCESSSEPGLPAQGCVYELPKRNL
RSSDAGTYYCAVATCGQILFGNGTRLEF

**>4857_c0_g1_i2**
MSSSLPDSLCFPGLLLCQLFAQSPSSVTVELGGSVTLSCSVSKSYSSNVLVWIKQPLGKLSISIASFQHNEASFQFEFYNNKRFAIDRRGDYFNLKISNIEA
SDVARYYCGAIRDTTVRFGSGTSIRLKGSGSVSRRVEQQPASVPVQPGDSVTLQCTIHTETCAGEHSVHWFRHGSGEALPGLIYTHGNRSDPCESSSEPGLP
AQGCVYELPKRNLRSSDAGTYYCAVATCGQILFGNGTRLEF

**>4857_c0_g1_i4**
MNIILFLLSFCSKAGLLLCQLFAQSPSSVTVELGGSVTLSCSVSKSYSSNALVWIKQPSGEPSTSIVSFKNNAALFHGGFDNNKRCAIDRSGDSFNLNISNI
EASDVARYYCGAIRDTTVRFGSGTSIRLKGSGSVSRRVEQQPASVPVQPGDSVTLQCTIHTETCAGEHSVHWFRHGSGEALPGLIYTHGNRSDPCESSSEPG
LPAQGCVYELPKRNLRSSDAGTYYCAVATCGQILFGNGTRLEF

**>6486_c3_g1_i3**
ALVWIKQPSGGPSTSIVSFKNNAALFHGGFDNNKRCAIDRSGDSFNLNISNIEASDVARYYCGAIRDTTVRFGSGTSIRLKGSGSVSRRVEQQPASVPVQPG
DSVTLQCTIHTETCAGEHSVHWFRHGSGEALPGLIYTHGNRSDPCESSSEPGLPAQGCVYELPKRNLRSSDAGTYYCAVATCGQILFGNGTRLHIAGDRSEA
GGSAEGPCGCGGHSALSLSVLALAAVLCVFGTLVTALLCRRHRAGTGPQGAEAASGRPPHSDQHCSPAHGTDTESQRGERDV**LAYAAL**DFDQNKRKAGKKRR
EG

**>6505_c0_g1_i10**
ALVWIKQPSGGPSTSIVSFKNNAALFHGGFDNNKRCAIDRRGDYFNLKISNIEASDVARYYCGGIRETSVGFGSGTTIRLKGSFSRTIVQWPVSVFVQPGDS
VTLQCTIHTETCAGEHSVHWFRHGSGEALPGLIYTHGNRSDPCESSSEPGLPAQGCVYELPKRNLRSSDAGTYYCAVSTCGQILFGNGTRLDIAGQTSLFPL
VLALGVSNAVCVLVIVVLIYTRNTRCAGDSQSSPDAVNTHTSHQNQDTEL**LNYAAL**KFTANTAVHPGRR

**>6505_c0_g1_i12**
MIRLCVTLLLFCKVYVNEGTVRSTVLRTAQLGDSVTLLCDLQGSYLIWFKQTTAQTPRSIATSYNYLPEITLFNEFKTDARFTVKRDEEGFNLTIARTEPSD
EATYFCGILRTNHVHFGNGTYLTLKGSKSVSRRVEQQPASVFVQPGDSVTLQCTIHTETCAGEHSVHWFRHGSGEALPGLIYTHGNRSDPCESSSEPGLPAQ
GCVYELPKRNLRSSDAGTYYCAVSTCGQILFGNGTWLDIAGKQCSAAELTNAQNQNS

**>6505_c0_g1_i13**
ALVWIKQPSGGPSTSIVSFKNNAALFHGGFDNNKRCAIDRRGDYFNLKISNIEASDVARYYCGGIRETSVGFGSGTTIRLKGSFSRTIVQWPVSVFVQPGDS
VTLQCTIHTETCAGEHSVHWFRHGSGEALPGLIYTHGNRSDPCESSSEPGLPAQGCVYELPKRNLRSSDAGTYYCAVSTCGQILFGNGTRLEFAGDRSEAGG
SAEGPSGCGGHSALSLSVLALAAVLCVFGTLVTALLCRRHRAGTGPQGAEAASGRPPHSDQHCSPAHGTDTESQQRGE*

**>6505_c0_g1_i4**
MIRLCVTLLLFCQVYLAQTENVVQPRLYTAAQLGDSVTLECFLPSQKSTYTVWFKQTIGQRPQCMATAYNYLQETTFYEEFKHNPRFTVQRERDSFHLNISR
TELSDTATYYCGVMFLNHMQFGNGTFLMVKGSVSRRVEQQPASVFVQPGDSVTLQCTIHTETCAGEHSVHWFRHGSGEALPGLIYTHGNRSDPCESSSEPGL
PAQGCVYELPKRNLRSSDAGTYYCAVSTCGQILFGNGTRLEFAGDRSEAGGSAEGPSGCGGHSALSLSVLALAAVLCVFGTLVTALLCRRHRAGTGPQGAEA
ASGRPPHSDQHCSPAHGTDTESQRGE*

**>6505_c0_g1_i8**
MIRLCVMLLLFCKVYLVQTEDVVQPRLFTAAQLGDSVTLECFISTERINYLSWIKQTVGQRPRGIVTSYSYLEDFTFYDEFRNNPRFTVQKDKGLFHLNISR
AEQSDAATYYCVTIYQNSVQFGNGTVLMLKGLESNMTIVQQPASVPVQPG

**>6505_c0_g1_i9**
MITLRLTLAILWTNYVFPEVVVSQPRLSVSARLGDPVTLQCYTGVEMVKTWMWLKHTIGQAPRTVITTYYDKIELYGDFNNTRVIAKKDNGNFTVTFSHTKS
LDVATYYCGIQLYSLYHYGNGTFVMLKGSKSVIRRVEQQPASVFVQPGDSVTLQCTIHTETCAGEHSVHWFRHGSGEALPGLIYTHGNRSDPCESSSEPGLP
AQGCVYELPKRNLRSSDAGTYYCAVSTCGQILFGNGTRLEFAGDRSEAGGSAEGPSGCGGHSALSLSVLALAAVLCVFGTLVTALLCRRHRAGTGPQGAEAA
SGRPPHSDQHCSPAHGTDTESQRGE*

**>6505_c13_g1_i1**
MITPGLTLAVLWMYCVNSTLLSQPHLSVSVRLGDPVTLECVATSDRGTRAWLKQSVGEAPSSIVTSYYETDKFHGRFKDDNRLTVQKDKTSFNLTFSRTESS
DAAAYFCGIIAYNSIDFGKGTFLVIIGSESVNRTVEQQPAS

**>6505_c17_g1_i1**
ECYINSETFSYVSWFKQSIGQRPHPIAVIYARLSDAMFLDEFKDNPRFKVYKEDTHFNLTISRTETSDTATYYCGKSYINVVHFGSGTFLLVTGSESTSRRV
EQQ

**>6505_c28_g1_i1**
HVFSHDGFVSDMTQTAATTLPFFFKTVQRGDSVTLGCLLATDRISFKAWFKQSTGFIPQVIAYSDMYLEKATFHNEFKDDPRFAVKADNHFQLTISGAEPSD
SAAYYCGSLYLKKWQFGNGTFVMVRESESVIRIVEQQPASVPVQP

**>6505_c30_g1_i1**
DSVTLECYVPSDKVSYMSWFKQTIGQKPRPIAVSYAHVSDATFLDEFKGNPRFKVHTEDTQYHLTISRTEPSDTATYY

**>6505_c3_g1_i7**
MIRLCLPLCLLRTSVILADAITQPGFLLRAQLGDSVSLQCFIHQQSLKQLFWYKQVVGEAPKCMFSSYGQSGQLTRHGEFNHSRFSVQRTGSAFHLCISNTE
PTDVGTYYCAAHDCSQVLFGGGSTLLLRDTDSRFIQPPLFQSLQSGLNSDLQCMMESRNLSADHRVHWFIKLSGQPSSKTPFTAGSMIQSSSEDDLPAQGCV
YNLSKRNLSCSDAESVYCAVATCGEILFGKGTEQEVTGRGVGVSHVPVVLLGALNAVLVVVIAVLLCRRQQSAQCQCCAARESGPVSRSPPQHSTSLQNETI
PT**VTYAAL**DFHHIKSKKERSKSVKKD**TVYSDI**MHQRWE*

**>7779_c0_g1_i4**
HTETCAGEHSVHWFRHGSGEALPGLIYTHGNRSDPCESSSEPGLPAQGCVYELPKRNLRSSDAGTYYCAVATCGQILFGNGTRLEFAGDQAEPLEPLERPCA
AQLSVLAWLSVARTAALLLCVALLAVIYCRGPER*

**>9858_c0_g1_i1**
LGSKWVSRSVVQQLASVPVQPGDSVTLQCTIHTETCAGEHSVHWFRHGSGEALPGLIYTHGNRSDPCESSSEPGLPAQGCVYELPMRNLRSSDAGTYYCAVA
TCGQILFGNGT

**>9872_c0_g1_i1**
RSSDVAVYYCAAFMYRRVACGNGTLLLLKVSGSVSRRVEQQPASVPVQPGDSVTLQCTIHTETCAGEHSVHWFRHGSGEALPGLIYTHGNRSDPCESSSEPG
LPAQGCVYELPKRNLRSSDAGTYYCAVATCGQILFGNGTRLEFAG

**Fig. S11. Transcriptome-predicted bowfin NITR proteins.**

NITR V domains are shaded dark blue and I domains are shaded light blue. Signal peptides and transmembrane (TM) domains are boxed. ITIM and ITIM-like sequences are in red text and a charged residue within a TM domain are shaded green.

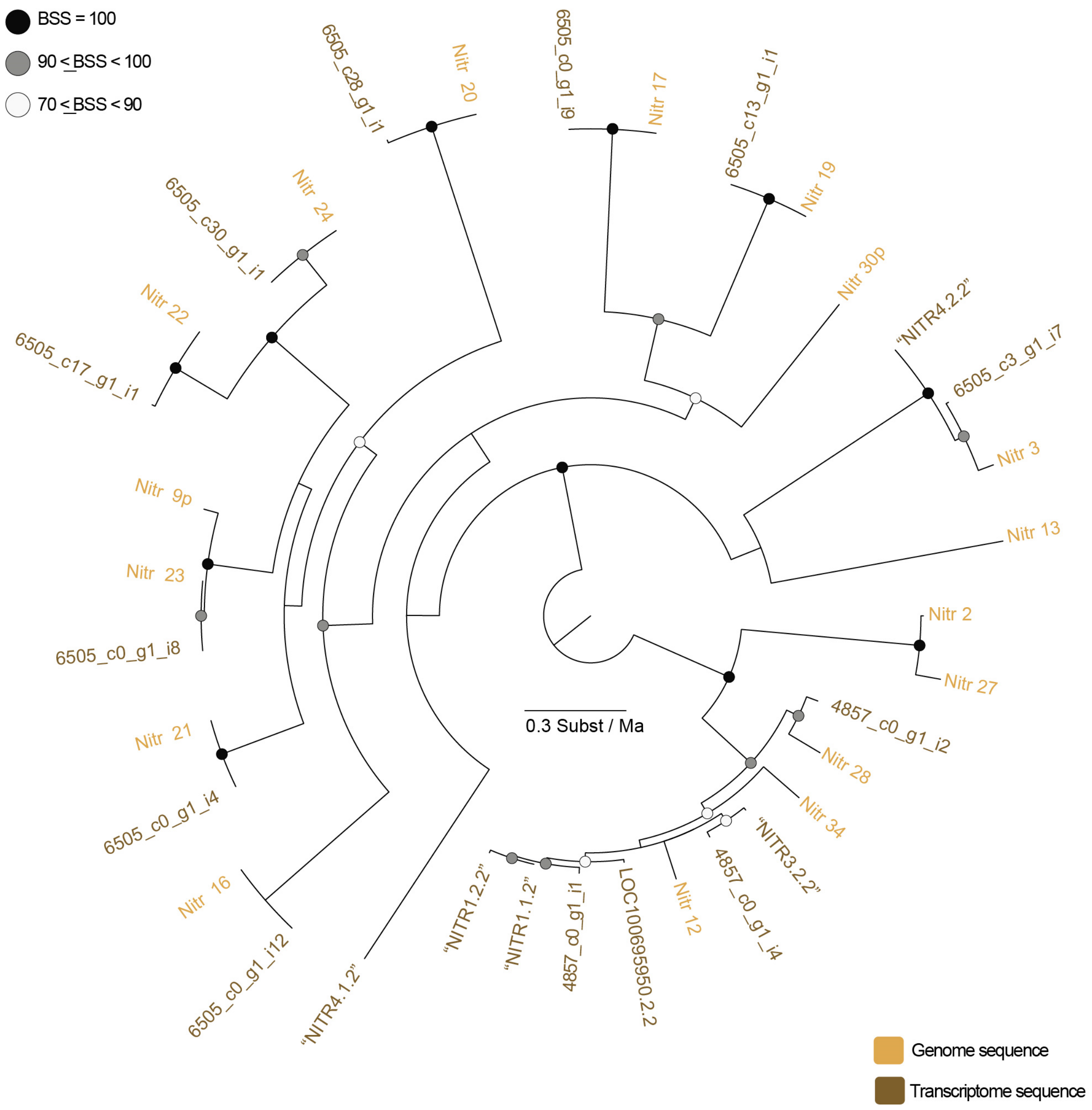

**Fig. S12. Bowfin NITR transcripts do not match (exactly) the reference genome.**
The phylogenetic relationships of NITR V domains identified from the bowfin genome (light brown) and transcripts (dark brown) inferred using maximum likelihood in IQ-TREE. Circles at nodes indicate bootstrap support values (BSS) with filled black circles black indicating BSS=100, gray circles indicating BSS values equal to or greater than 90 but less than 100, and white circles indicating BSS values greater than 70 but less than 90.

```
Nitr16             1 MIRLCVTLLLFCKV-VNEGTVRSTVLRTAQLGDSVTLLCDLQGSYLIWFKQTTAQTPRSIATSYNYLPEITLFNEFKTDA
6505_c0_g1_i12     1 MIRLCVTLLLFCKVYVNEGTVRSTVLRTAQLGDSVTLLCDLQGSYLIWFKQTTAQTPRSIATSYNYLPEITLFNEFKTDA

Nitr16            80 RFTVKRDEEGFNLTIARTEPSDEATYFCGILRTNHVHFGNGTYLTLKSGSKSVSRRVEQQPASVPVQSGDSVTLQCTIYT
6505_c0_g1_i12    81 RFTVKRDEEGFNLTIARTEPSDEATYFCGILRTNHVHFGNGTYLTLK-GSKSVSRRVEQQPASVFVQPGDSVTLQCTIHT

Nitr16           160 EPCAGEHSVHWFRHGSGEALPGLIYTHGNRSDPCESSSEPGLPAQGCVYELPKRNLCSSDAGTYYCAVATCGQILFGNGT
6505_c0_g1_i12   160 ETCAGEHSVHWFRHGSGEALPGLIYTHGNRSDPCESSSEPGLPAQGCVYELPKRNLRSSDAGTYYCAVSTCGQILFGNGT

Nitr16           240 RMDIA-----------------
6505_c0_g1_i12   240 WLDIAGKQCSAAELTNAQNQNS
```

**Fig. S13. Nitr16 and 6505_c0_g1_i12.**

Alignment of bowfin proteins encoded by the *nitr16* gene and *6505_c0_g1_i12* transcript.

```
Nitr17             1 MITLRLTLAILWTN-VFPEVVVSQPRLSVSARLGDPVTLQCYTGVEMVKTWMWLKHTIGQAPRTVITTYYDKIELYGDFN
6505_c0_g1_i9      1 MITLRLTLAILWTNYVFPEVVVSQPRLSVSARLGDPVTLQCYTGVEMVKTWMWLKHTIGQAPRTVITTYYDKIELYGDFN

Nitr17            80 NTRVIAKKDNGNFTVTFSHTKSLDVATYYCGIQLYSLYHYGNGTFVMLK-------------------------------
6505_c0_g1_i9     81 NTRVIAKKDNGNFTVTFSHTKSLDVATYYCGIQLYSLYHYGNGTFVMLKGSKSVIRRVEQQPASVFVQPGDSVTLQCTIH

Nitr17               --------------------------------------------------------------------------------
6505_c0_g1_i9    161 TETCAGEHSVHWFRHGSGEALPGLIYTHGNRSDPCESSSEPGLPAQGCVYELPKRNLRSSDAGTYYCAVSTCGQILFGNG

Nitr17               --------------------------------------------------------------------------------
6505_c0_g1_i9    241 TRLEFAGDRSEAGGSAEGPSGCGGHSALSLSVLALAAVLCVFGTLVTALLCRRHRAGTGPQGAEAASGRPPHSDQHCSPA

Nitr17               ------------
6505_c0_g1_i9    321 HGTDTESQRGE*
```

**Fig. S14. Nitr17 and 6505_c0_g1_i9.**

Alignment of bowfin proteins encoded by the *nitr17* gene and *6505_c0_g1_i9* transcript.

```
Nitr21            1 MIRLCVTLLLFCQV-LAQTENVVQPRLYTAAQLGDSVTLECFLPSQKSTYTVWFKQTIGQRPQCMATAYNYLQETTFYEE
6505_c0_g1_i4     1 MIRLCVTLLLFCQVYLAQTENVVQPRLYTAAQLGDSVTLECFLPSQKSTYTVWFKQTIGQRPQCMATAYNYLQETTFYEE

Nitr21           80 FKHNPRFTVQRERDSFHLNISRTELSDTATYYCGVMFLNHMQFGNGTFLMVKISGSVSRRVEQQPASVPVQPGDSVTLQC
6505_c0_g1_i4    81 FKHNPRFTVQRERDSFHLNISRTELSDTATYYCGVMFLNHMQFGNGTFLMVK--GSVSRRVEQQPASVFVQPGDSVTLQC

Nitr21          160 TIHTETCAGEHSVHWFRHGSGEALPGLIYTHGNRSDPCESSSEPGLPAQGCVYELPKRNLRSSDAGTYYCAVATCGQILF
6505_c0_g1_i4   159 TIHTETCAGEHSVHWFRHGSGEALPGLIYTHGNRSDPCESSSEPGLPAQGCVYELPKRNLRSSDAGTYYCAVSTCGQILF

Nitr21          240 GNGTRLEFAG----------------------------------------------------------------------
6505_c0_g1_i4   239 GNGTRLEFAGDRSEAGGSAEGPSGCGGHSALSLSVLALAAVLCVFGTLVTALLCRRHRAGTGPQGAEAASGRPPHSDQHC

Nitr21              ---------------
6505_c0_g1_i4   319 SPAHGTDTESQRGE*
```

**Fig. S15. Nitr21 and 6505_c0_g1_i4.**

Alignment of bowfin proteins encoded by the *nitr21* gene and *6505_c0_g1_i4* transcript.
